# Towards Effective CAIX-targeted Radionuclide and Checkpoint Inhibition Combination Therapy for Advanced Clear Cell Renal Cell Carcinoma

**DOI:** 10.1101/2024.02.16.580614

**Authors:** Simone C. Kleinendorst, Egbert Oosterwijk, Janneke Molkenboer-Kuenen, Cathelijne Frielink, Gerben M. Franssen, Daan F. Boreel, Giulia Tamborino, Manon Gloudemans, Merel Hendrikx, Dennis Kroon, Jopp Hillen, Johan Bussink, Stijn Muselaers, Peter Mulders, Mark W. Konijnenberg, Michael P. Wheatcroft, Kwame Twumasi-Boateng, Sandra Heskamp

## Abstract

**Background:** Immune checkpoint inhibitors (ICI) are routinely used in advanced clear cell renal cell carcinoma (ccRCC). However, a substantial group of patients does not respond to ICI therapy. Radiation is a promising approach to increase ICI response rates since it can generate anti-tumor immunity. Targeted radionuclide therapy (TRT) is a systemic radiation treatment, ideally suited for precision irradiation of metastasized cancer. Therefore, the aim of this study is to explore the potential of combined TRT, targeting carbonic anhydrase IX (CAIX) which is overexpressed in ccRCC, using [^177^Lu]Lu-DOTA-hG250, and ICI for the treatment of ccRCC.

**Methods:** In this study, we evaluated the therapeutic and immunological action of [^177^Lu]Lu-DOTA-hG250 combined with aPD-1/a-CTLA-4 ICI. First, the biodistribution of [^177^Lu]Lu-DOTA-hG250 was investigated in BALB/cAnNRj mice bearing Renca-CAIX or CT26-CAIX tumors. Renca-CAIX and CT26-CAIX tumors are characterized by poor versus extensive T-cell infiltration and homogeneous versus heterogeneous PD-L1 expression, respectively. Tumor-absorbed radiation doses were estimated through dosimetry. Subsequently, [^177^Lu]Lu-DOTA-hG250 TRT efficacy with and without ICI was evaluated by monitoring tumor growth and survival. Therapy-induced changes in the tumor microenvironment were studied by collection of tumor tissue before and 5 or 8 days after treatment and analyzed by immunohistochemistry, flow cytometry, and RNA profiling.

**Results:** Biodistribution studies showed high tumor uptake of [^177^Lu]Lu-DOTA-hG250 in both tumor models. Dose escalation therapy studies in Renca-CAIX tumor-bearing mice demonstrated dose-dependent anti-tumor efficacy of [^177^Lu]Lu-DOTA-hG250 and remarkable therapeutic synergy including complete remissions when a presumed subtherapeutic TRT dose (4 MBq, which had no significant efficacy as monotherapy) was combined with aPD-1+aCTLA-4. Similar results were obtained in the CT26-CAIX model for 4 MBq [^177^Lu]Lu-DOTA-hG250 + a-PD1. *Ex vivo* analyses of treated tumors revealed DNA damage, T-cell infiltration, and modulated immune signaling pathways in the TME after combination treatment.

**Conclusions:** Subtherapeutic [^177^Lu]Lu-DOTA-hG250 combined with ICI showed superior therapeutic outcome and significantly altered the TME. Our results underline the importance of investigating this combination treatment for patients with advanced ccRCC in a clinical setting. Further investigations should focus on how the combination therapy should be optimally applied in the future.

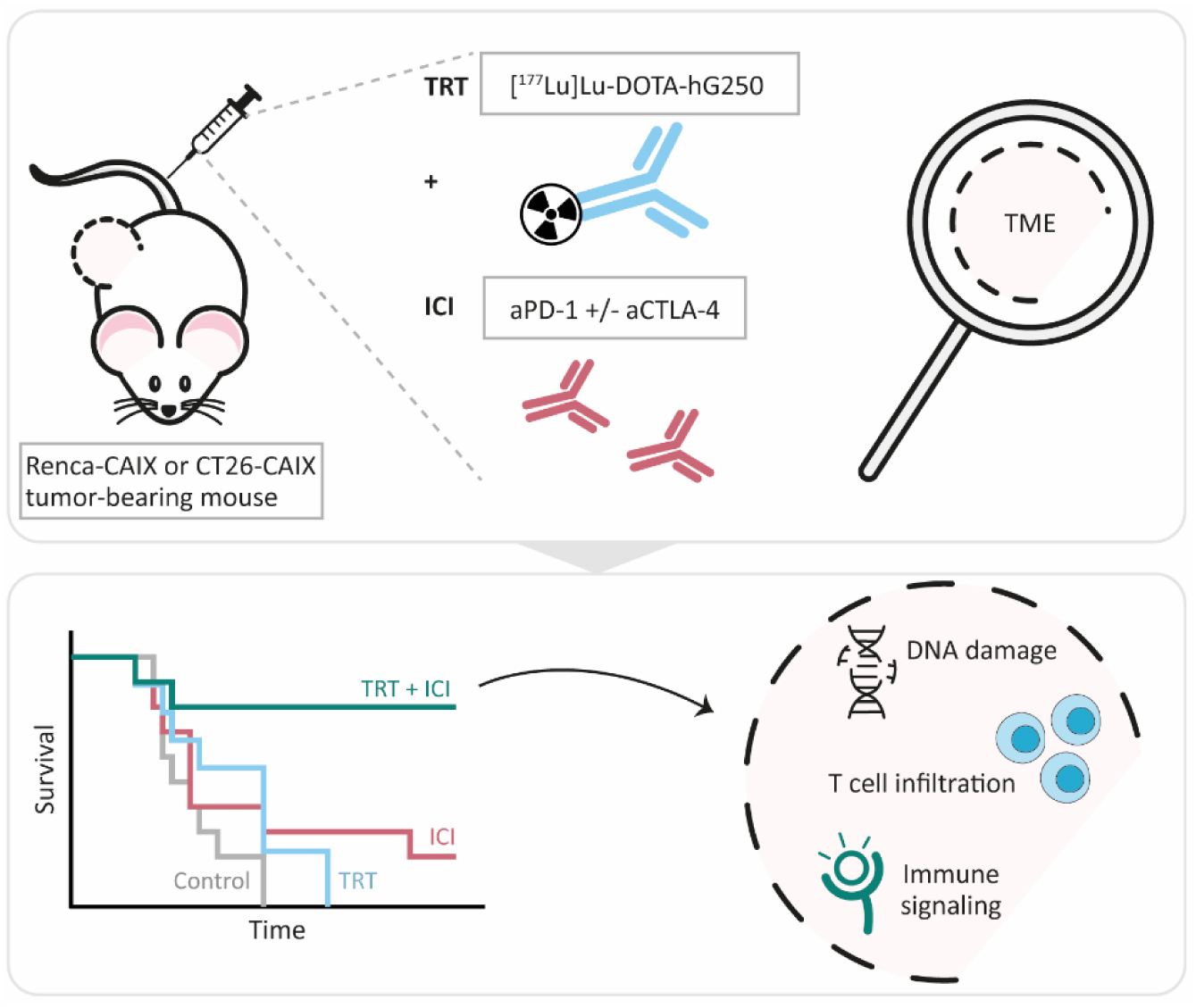

## BACKGROUND

Clear cell renal cell carcinoma (ccRCC) is the most common subtype of kidney cancer, being >85% of metastatic RCC cases [1]. Treatment of advanced ccRCC has evolved substantially with the implementation of tyrosine kinase inhibitors (TKIs) and immune checkpoint inhibitors (ICIs) as a standard of care according to the EAU guideline [2]. However, a significant portion of RCC patients does not respond to these treatment strategies, which underlines the ongoing clinical need for novel therapeutic approaches resulting in effective and durable responses [3].

A widely studied combination regimen is ICI with external beam radiation therapy (EBRT), in which the central approach is to overcome ICI resistance by driving the tumor microenvironment (TME) towards a more immunogenic phenotype [4, 5]. In addition to its direct effects on DNA, radiation can trigger a cascade of immunological events. For example, the induction of immunogenic cell death via the release of danger-associated molecular patterns such as adenosine triphosphate (ATP), high-mobility group box 1 (HMGB1), and calreticulin [6]. Furthermore, radiation can generate neo-antigens, upregulate major histocompatibility (MHC) molecules, enhance cytokine release, and activate the type I interferon (IFN) pathway, together resulting in increased immune cell recruitment and activation [4, 5, 7]. However, local irradiation with EBRT might be insufficient to generate a systemic anti-tumor immune response in patients with metastatic cancer, possibly also limited by immunosuppression in (un)irradiated lesions, as observed in clinical studies combining EBRT with immunotherapy [8].

An alternative to EBRT is targeted radionuclide therapy (TRT). With this systemic radiation treatment, a tumor-targeting radiopharmaceutical is injected intravenously resulting in selective radiotracer uptake and consequently irradiation of metastatic lesions [9]. A suitable radiotracer for the treatment of patients with advanced ccRCC is [^177^Lu]Lu-DOTA-girentuximab (G250) [10, 11]. This radiotracer specifically targets carbonic anhydrase IX (CAIX), a transmembrane protein overexpressed in >95% of ccRCCs, as a consequence of mutations in the Von Hippel Lindau (VHL) gene with limited expression in healthy tissue [12]. Phase-I and II clinical trials with [^177^Lu]Lu-DOTA-girentuximab have shown a clear indication of efficacy [10, 11]. However, in this selected patient population profound myelotoxicity was observed, which limited the radiation dose and the possibility for retreatment with TRT. This makes [^177^Lu]Lu-girentuximab currently only moderately effective as monotherapy.

Previous preclinical studies on combined TRT and ICI have shown promising results in several solid tumors such as melanoma and colon adenocarcinoma [13, 14]. Similarly, combination of CAIX-TRT with ICI may hold promise for the treatment of advanced ccRCC. However, the therapeutic responses in these TRT/ICI combination studies are diverse and the optimal design of the combined regimen, including radionuclide choice, dose, frequency, and timing, for synergy is not fully understood. Furthermore, our knowledge of how therapeutic action relates to tumor types, e.g. inflamed and desert phenotypes, is limited. Therefore, additional preclinical studies are needed to broaden our understanding of TRT and the accompanying radiobiology to aid the design of future clinical trials.

In the current study, we aim to assess the therapeutic efficacy of combined [^177^Lu]Lu-DOTA-hG250 and aPD-1/aCTLA-4 therapy in syngeneic mouse models and to explore its effects on the TME.

## METHODS

### Antibodies and cell lines

CAIX-directed humanized IgG1 monoclonal antibody girentuximab (hG250) was provided by Telix Pharmaceuticals (Melbourne, Australia). Anti-murine PD-1 and anti-murine CTLA-4 were purchased from Bio X Cell (BE0146-clone RMP1-14, and BE0032-clone UC10-4F10-11). SKRC-52, CT26 WT, CT26-CAIX, Renca WT, and Renca-CAIX cells (**Table S1**) were cultured in base medium (RPMI1640 (Gibco) supplemented with glutamine (2 mM, Gibco) and fetal calf serum (10%, FCS, Sigma-Aldrich-Chemie-BV)) with Geneticin G418 Sulphate (0.6 mg/ml, 11811-031, Gibco) for CAIX-transfected lines, and non-essential amino acids (0.1 mM Gibco) and sodium pyruvate (1 mM, Gibco) for Renca lines. Cells were cultured at 37°C in a humidified atmosphere with 5% CO2, and passaged maximally ten times for all experiments.

### CAIX expression and hG250 binding

The CAIX density on cells was determined by Scatchard analysis using [^111^In]In-DOTA-hG250 (0.37 MBq/µg). Cell lines were cultured to confluency in 6-well plates and subsequently incubated with binding buffer (RPMI1640, 0.5% bovine serum albumin) containing increasing concentrations of [^111^In]In-DOTA-hG250 (0.003 – 30 nM) for 4h on ice. Unbound [^111^In]In-DOTA-hG250 was removed and the cell-associated activity was measured in a γ-counter (1480 Wizard 3; LKB/Wallace, Perkin Elmer, Boston, MA, USA).

### Clonogenic survival and cell viability assays

Radiosensitivity to EBRT was determined using clonogenic survival assays. Cells were seeded as single cells, attached overnight, and irradiated with 0-8 Gy using the XRAD 320 (RPS Services Limited) at a dose rate of 3.1 Gy/min and cultured for another 9 days. Cells were fixed and stained with crystal violet (0.5% in 50% methanol, 20% ethanol, 30% water), and colonies (>50 cells) were manually scored under the microscope. Surviving fraction was determined by dividing the ratio of colonies and seeding density at x Gy by these values at 0 Gy and the survival data was described using the linear-quadratic model with weighted sum of squares (1/y^k, k=1.5).

### Animal experiments

All experiments were performed in accordance with the principles laid out by the revised Dutch Act on Animal Experimentation (1997) and a protocol for each experiment was approved by the Radboudumc institutional Animal Welfare Body. Animals were housed and fed according to Dutch animal welfare regulations. The experiments were performed in female BALB/cAnNRj mice (*n*=347, 10-12 weeks old, 20-25 gram, Janvier, le Genest-Saint-Isle, France). Mice were accustomed to laboratory conditions for 1 week and housed in individualized ventilated cages with *ad libitum* access to animal chow and water. To reduce distress from tumor growth, wet food was provided daily after intervention. All interventions were given in 0.9% NaCl at a volume of 200 µL. Renca-CAIX or CT26-CAIX tumors were engrafted by subcutaneous injection of 1·10^6^ or 5·10^5^ cells, respectively, in the right flank. Tumor take rate was approximately 83% and 71%, respectively and animals were block-randomized into treatment groups based on tumor volume when tumors reached a mean size of 50-100 mm^3^, as determined by caliper measurements performed thrice a week and calculated according to an ellipsoid model (with 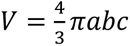 (a, b, and c being the tumor radii)). Animals were excluded from experiments if tumor size was not measurable at timepoint of treatment. Details on sample size calculations, dropouts, randomization, and blinding are described in supplemental methods. Animals were sacrificed by CO_2_/O_2_ asphyxiation at the end of study or when mice reached humane endpoints (tumor volume ≥ 1.5 cm^3^, tumor ulceration, >15% body weight loss within 2 d, >20% body weight loss compared with start weight, or severe clinical deterioration as assessed by a biotechnician).

### Biodistribution studies and dosimetry

Biodistribution of [^177^Lu]Lu-DOTA-hG250 (0.1 MBq/µg) was determined in Renca-CAIX- (*N*=4-5) or CT26-CAIX- (*N*=2-3) tumor-bearing mice. Mice received 0.2 MBq [^177^Lu]Lu-DOTA-hG250 at a protein dose of 3 µg via tail vein injection. Mice were sacrificed at 1, 3, and 7 days post injection and tumor and normal tissues were harvested, weighed, and activity was measured in a γ-counter. The uptake in all organs was expressed as percentage of injected activity per gram tissue (%IA/g). Absorbed dose estimations were performed using the time-activity curves for dosimetry as described in supplemental methods.

### Therapy studies

Therapeutic efficacy of [^177^Lu]Lu-DOTA-hG250 monotherapy at different doses was studied in Renca-CAIX tumor-bearing mice. Mice (*n*=10) were injected with 5 µg of 12, 18, or 24 MBq [^177^Lu]Lu-DOTA-hG250 or vehicle control (0.9% NaCl) via intravenous tail vein injection.

To assess therapeutic efficacy of combined TRT/ICI therapy, the following treatment regimens were administered to Renca-CAIX tumor-bearing mice (*n*=10): (1) vehicle, (2) TRT alone, (3) ICI alone, and (4) TRT/ICI. All groups received one intravenous injection on day 0 (0.9% NaCl for vehicle and ICI alone, and 4 or 12 MBq [^177^Lu]Lu-DOTA-hG250 for TRT alone and TRT/ICI) and eight intraperitoneal injections on day 1, 4, 7, 10, 13, 16, 19, and 22 (0.9% NaCl for vehicle and TRT alone, and 200 µg aPD-1 + 200 µg aCTLA-4 for ICI alone and TRT/ICI) (summarized in **Figure 1**).

**Figure 1.**
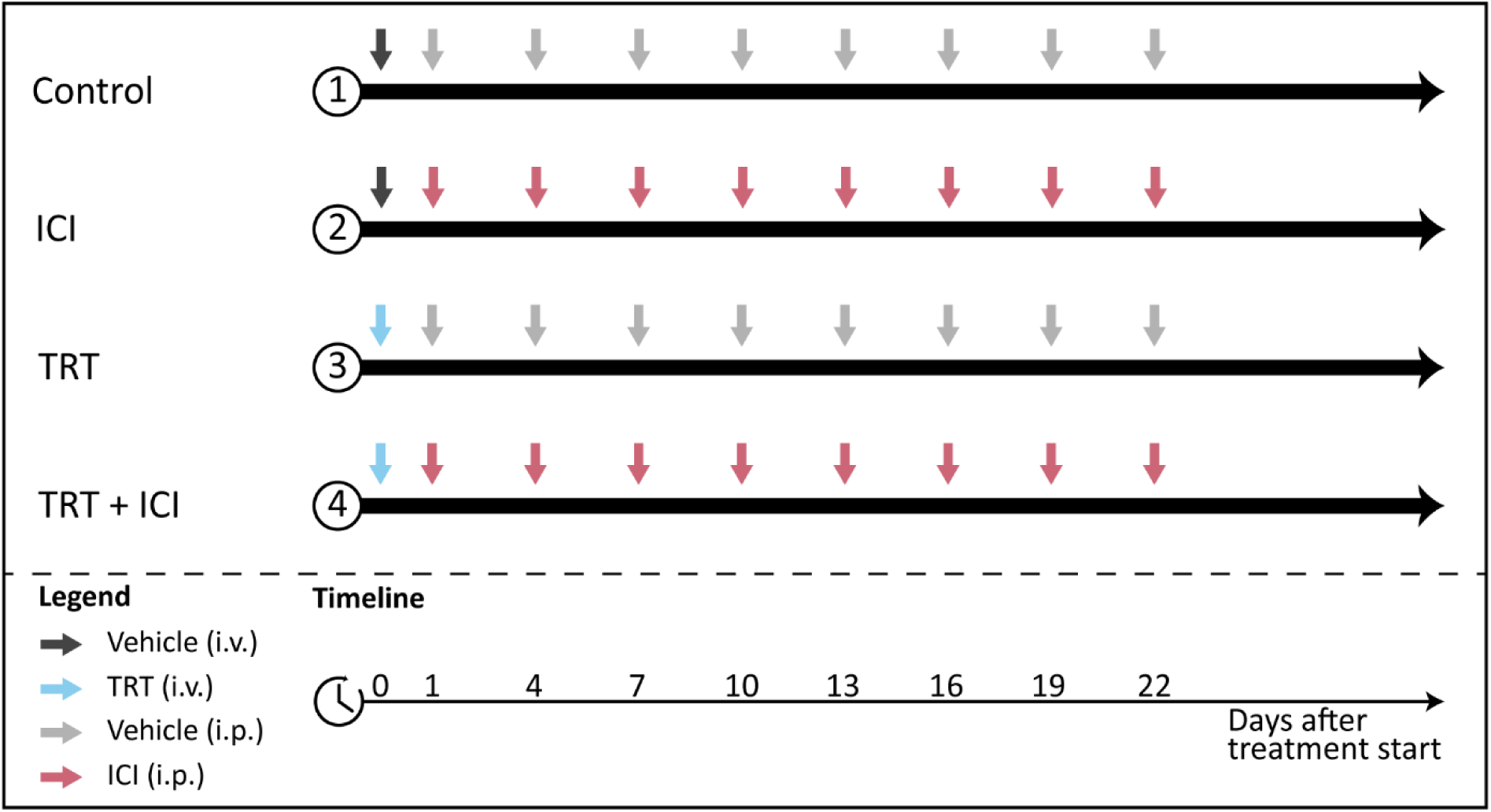
Combination therapy study design. Four treatment groups received different combinations of TRT (blue arrows) and ICI (pink arrows) or vehicle (grey arrows) injections.

For CT26-CAIX tumor-bearing mice (*n*=9-10) therapeutic efficacy of groups 1-4 was evaluated as described above, including only the 4 MBq TRT dose and ICI consisting of aPD-1 alone. Tumor volume and survival were monitored until the end of follow-up, predefined at 6 weeks post-treatment. Mice with complete tumor remission received a re-challenge tumor cell injection and were followed for another 4 weeks. The primary outcome of the study was tumor growth, which was expressed as normalized area under the curve (nAUC), calculated as area under tumor growth curves of single animals normalized to animal lifetime. Furthermore, percentage of mice with stable disease, defined as <10% fluctuation in tumor volume over ≥5 days, was determined using a tumor control index tool [15]. For the secondary outcome of the study, Kaplan-Meier survival curves were generated for each group.

Experiments for *ex vivo* analyses of the TME were performed twice, to collect whole fresh tumors for flow cytometry (part A) and to fix whole tumors in 4% formalin and embed in paraffin (FFPE) for immunohistochemistry and RNA profiling (part B). Renca-CAIX tumor-bearing mice (*n*=5-6) were sacrificed at day 0, or days 5 and 8 post-treatment (identical to the Renca-CAIX combination therapy experiment).

### Characterization of the tumor microenvironment

For flow cytometry analysis, fresh tumors were dissociated into single-cell suspension by incubating with collagenase III (1 mg/ml, Worthington, LS004182) and DNase I (0.1 mg/ml, Roche, 11284932001) in RPMI for 30 min at 37°C, and after adding 1 mM EDTA to stop the reaction passing the cells through a 100µm cell strainer (Corning, 431752) twice. Single cells were stained with antibody panels (**Table S2**).

For immunohistochemical analysis, FFPE tumor sections (4 µm) were evaluated for 53BP1, caspase-3, Ki67, CAIX, PD-L1, CD3, CD8, CD4, FOXP3, Ly6G, F4-80, and H&E, as described previously or in supplemental methods [16, 17]. Slides were digitized using a 3DHistech P1000 digital slide scanner (3DHistech, 20 × magnification, 0.24 µm/pixel) and four snapshots, selected manually to represent different viable tumor areas, per tumor were taken with CaseViewer 2.3. Fluorescent images were procured using a DM6000 fluorescence microscope (Leica). Images were quantified using ImageJ.

For RNA analysis, RNA was isolated from 10µm-thick sections of FFPE material using the RNeasy FFPE kit according to manufacturer’s instructions (Qiagen, 73504). RNA concentration was determined using a Nanodrop spectrophotometer (ThermoScientific) and RNA quality was determined using an Agilent Bioanalyzer. Expression of the 770 genes of the murine PanCancer IO 360 Gene Expression Panel (NanoString Technologies, Inc.) was quantified with the NanoString nCounter XT CodeSet Gene Expression Assay according to manufacturer’s instructions (NanoString Technologies, Inc., performed at St. Michael’s Hospital, Unity Health Toronto). For this, 100 ng of RNA was used per sample and samples were batch-hybridized for 21 hours.

Serum HMGB1 levels were determined using the HMGB1 express ELISA kit according to manufacturer’s instructions (Tecan, 30164033).

### Statistical analyses

CAIX expression was compared between cell lines using one-way ANOVA with Šidák multiple comparisons test. For clonogenic survival, the LQ model alpha/beta ratios were compared between the cell lines using one-way ANOVA with Tukey’s multiple comparison correction. For animal biodistribution and dosimetry studies, descriptive statistics were used. In animal therapy studies, all analyses entailed comparison of all treatment groups versus control group and combination treatment groups versus respective monotherapy groups. Differences in tumor growth (nAUC) between groups was compared using Kruskal-Wallis test with Dunn’s multiple comparisons test. Kaplan-Meier survival between the groups was compared using pairwise Mantel-Cox log-rank testing and Bonferroni correction. Differences in marker-positive areas (IHC) and immune cell abundance or marker expression (flow cytometry) were assessed using a mixed-effects model. RNA data analysis was performed using the Rosalind Platform (NanoString Technologies, Inc.), including pairwise comparisons of treatment groups to control and combination therapy to ICI, including timepoint as covariate and using cut-off values of minimally 1.5 or -1.5 fold change and Benjamini-Hochberg-adjusted p<0.05. Statistical significance was defined as p-value below 0.05, two-sided, and analyses were performed with Graphpad Prism, version 5.03.

## RESULTS

### Renca-CAIX and CT26-CAIX cells exhibit prominent *in vitro* CAIX expression and intrinsic radiosensitivity

CAIX expression of Renca-CAIX and CT26-CAIX was approximately 3-fold lower compared with the human ccRCC cell line SKRC-52, which has been previously used in *in vivo* studies assessing CAIX-TRT efficacy [18] (**Figure 2A, Figure S3**). Radiosensitivity of the cells to EBRT (**Figure 2B**) and [^177^Lu]Lu-DOTA-hG250 (**Figure S4**) was determined. The human SKRC-52 cell line was more radiosensitive than Renca-CAIX and CT26-CAIX cells to both EBRT and TRT, while no difference in radiosensitivity between Renca-CAIX and CT26-CAIX cell lines was observed (**Table S3**). Taken together, CAIX expression, hG250 binding, and intrinsic cell radiosensitivity were comparable between Renca-CAIX and CT26-CAIX cell lines but were lower for the SKRC-52 cell line.

**Figure 2.**
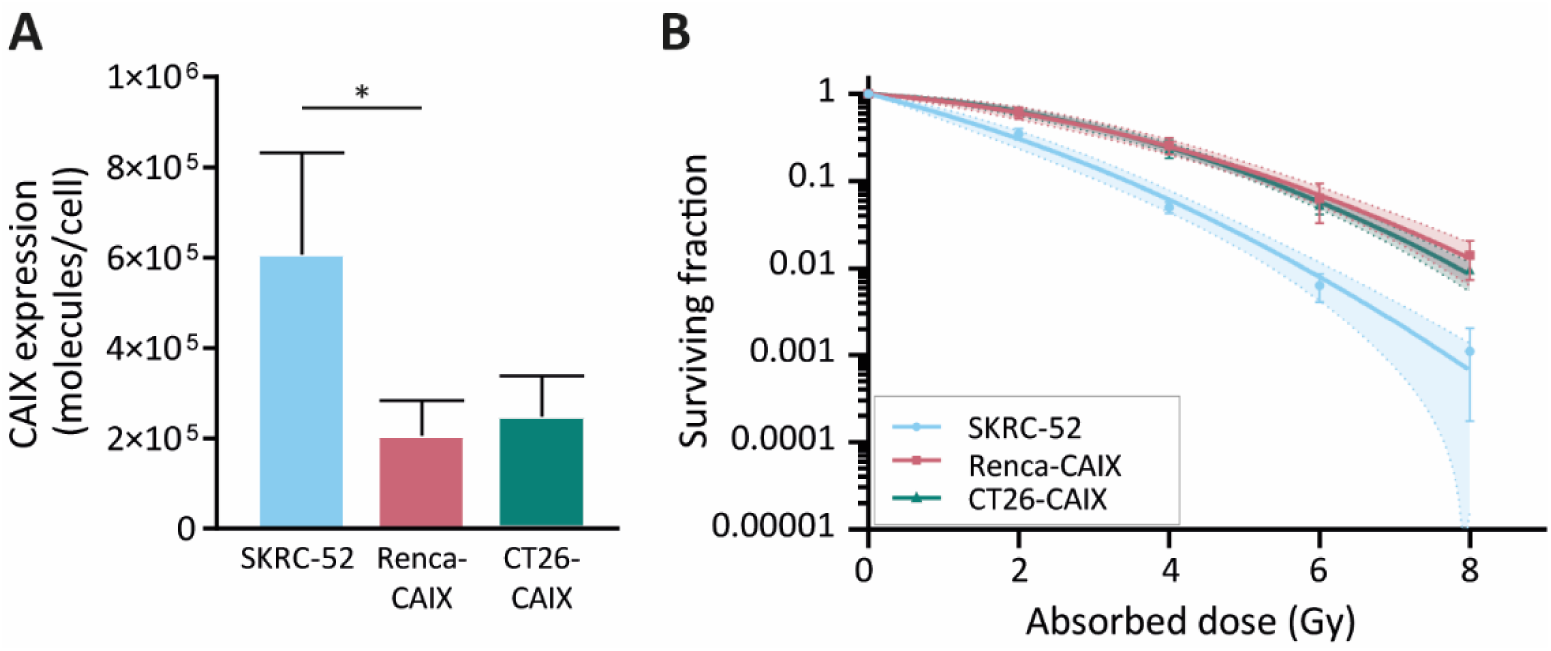
In vitro characterization of Renca-CAIX, and CT26-CAIX cell lines. **(A)** CAIX expression as determined by Scatchard analysis using [^111^In]In-DOTA-hG250. Data represent mean + SD of 4 independent experiments (* p<0.05). **(B)** Clonogenic survival of cells after radiation with 0-8 Gy EBRT. Data represent mean ± SD of 2 independent experiments. Non-linear regression using the linear quadratic model was used to fit the data and 95% confidence bands are shown (dashed lines).

### High *in vivo* Renca-CAIX and CT26-CAIX tumor uptake of [^177^Lu]Lu-DOTA-hG250

The biodistribution and pharmacokinetics of [^177^Lu]Lu-DOTA-hG250 were studied *ex vivo*, after dissection of Renca-CAIX and CT26-CAIX tumor-bearing mice at 1, 3, and 7 days post-treatment (**Figure 3A, Table S4**). For Renca-CAIX, the maximum tumor uptake was 32±9.4 %IA/g after 1 day which gradually decreased over time, while the maximum tumor uptake in CT26-CAIX was 17±2.3 %IA/g which remained stable over 7 days. [^177^Lu]Lu-DOTA-hG250 was cleared from blood over time, with only prominent off-target accumulation in liver and spleen. Dose estimations reported comparable tumor-absorbed doses for both tumor models (Renca-CAIX: 2.4±1.3 Gy/MBq, CT26-CAIX: 2.2±1.0 Gy/MBq). Liver-absorbed doses were estimated at 4.4±2.8 Gy/MBq and 3.3±0.9 Gy/MBq for Renca-CAIX and CT26-CAIX tumor-bearing mice, respectively. Furthermore, immunohistochemistry (IHC) showed abundant, but heterogeneous CAIX expression in both tumors, verifying that CAIX is expressed *in vivo* (**Figure 3B**). Generally, Renca-CAIX was characterized by prominent homogeneous PD-L1 expression and low abundance of T cells in the TME, while CT26-CAIX demonstrated heterogeneous PD-L1 expression throughout the tumor and was highly infiltrated by T cells. In summary, Renca-CAIX and CT26-CAIX tumor models demonstrated comparable biodistribution profiles of [^177^Lu]Lu-DOTA-hG250 but exhibited distinct immunologic characteristics.

**Figure 3.**
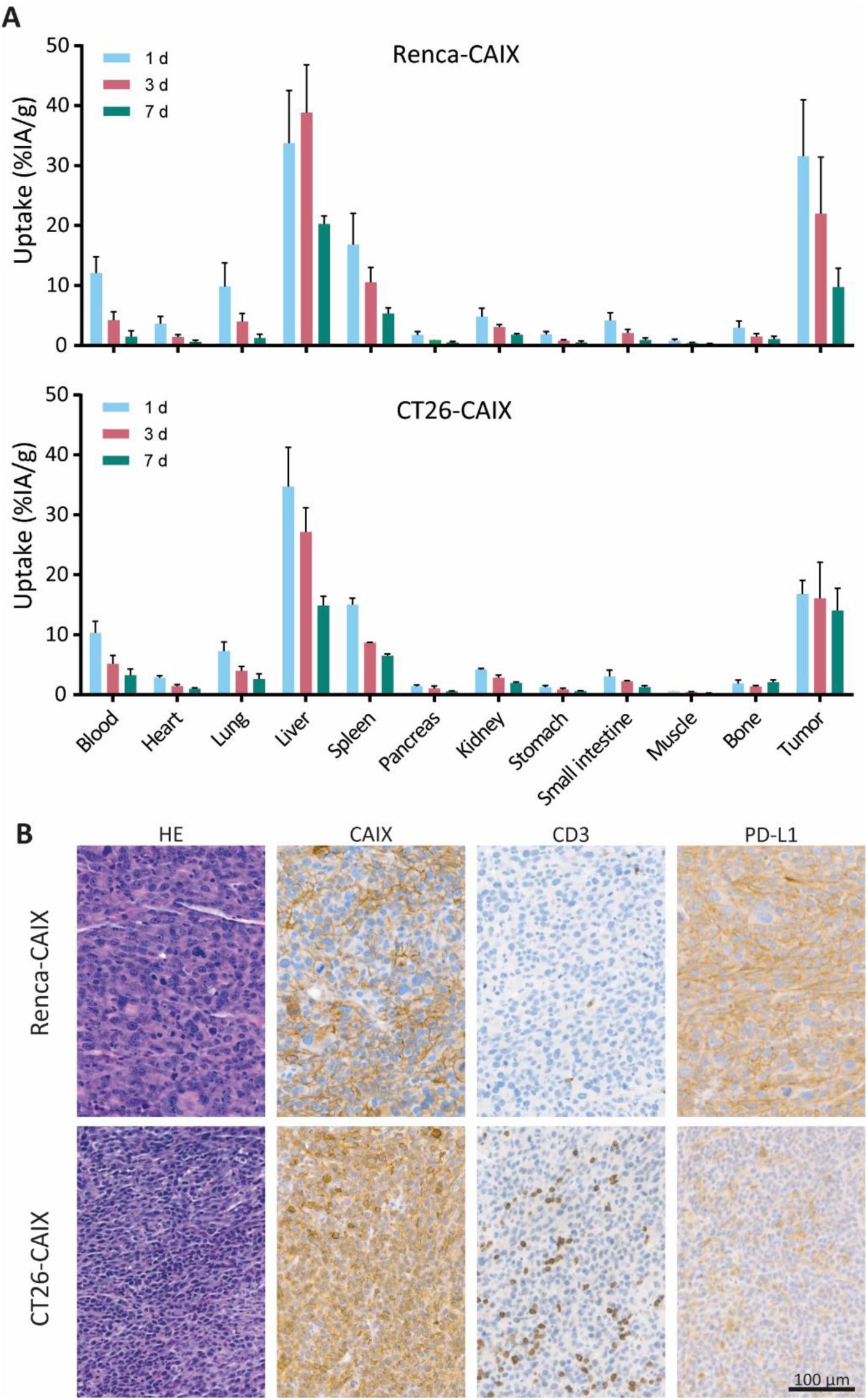
In vivo characterization of Renca-CAIX and CT26-CAIX tumor models. **(A)** Biodistribution of [^177^Lu]Lu-DOTA-hG250 (0.2 MBq / 3µg / mouse) as determined by dissection of mice at 1, 3, or 7 days post-treatment for Renca-CAIX (n = 5, 5, 4) and CT26-CAIX (n = 3, 2, 3). Data represents mean + SD. **(B)** Immunohistochemistry of untreated Renca-CAIX and CT26-CAIX tumors. Cell surface expression of CAIX, CD3, and PD-L1 are shown.

### [^177^Lu]Lu-DOTA-hG250 inhibits Renca-CAIX tumor growth and improves survival

Based on dosimetry, the liver was expected to be the dose-limiting organ for therapeutic studies, therefore a maximum injected activity of 24 MBq [^177^Lu]Lu-DOTA-hG250, resulting in a dose of 106±67 Gy to the liver for Renca-CAIX tumor-bearing mice, was used. Renca-CAIX tumor-bearing mice were randomized to receive 12, 18, or 24 MBq [^177^Lu]Lu-DOTA-hG250 (with estimated tumor-absorbed doses of 29, 44, and 59 Gy, respectively), or vehicle. Measurements of body weight and ALAT/ASAT liver enzymes gave no indication of toxicity (**Figure S5** and data not shown). Treatment with 18 or 24 MBq TRT significantly inhibited tumor growth compared to non-treated animals (p=0.03 and p<0.001, respectively) (**Figure 4A, Table S5**). Furthermore, these two treatments resulted in disease stabilization and complete tumor response in some animals, and significantly improved survival compared to non-treated animals (p=0.001, p=0.004, respectively) (**Figure 4B, Table S5**). Based on these results, to explore potential synergism or additive effects with ICI, subtherapeutic TRT activities of ≤12 MBq were chosen for subsequent combination therapy studies.

**Figure 4.**
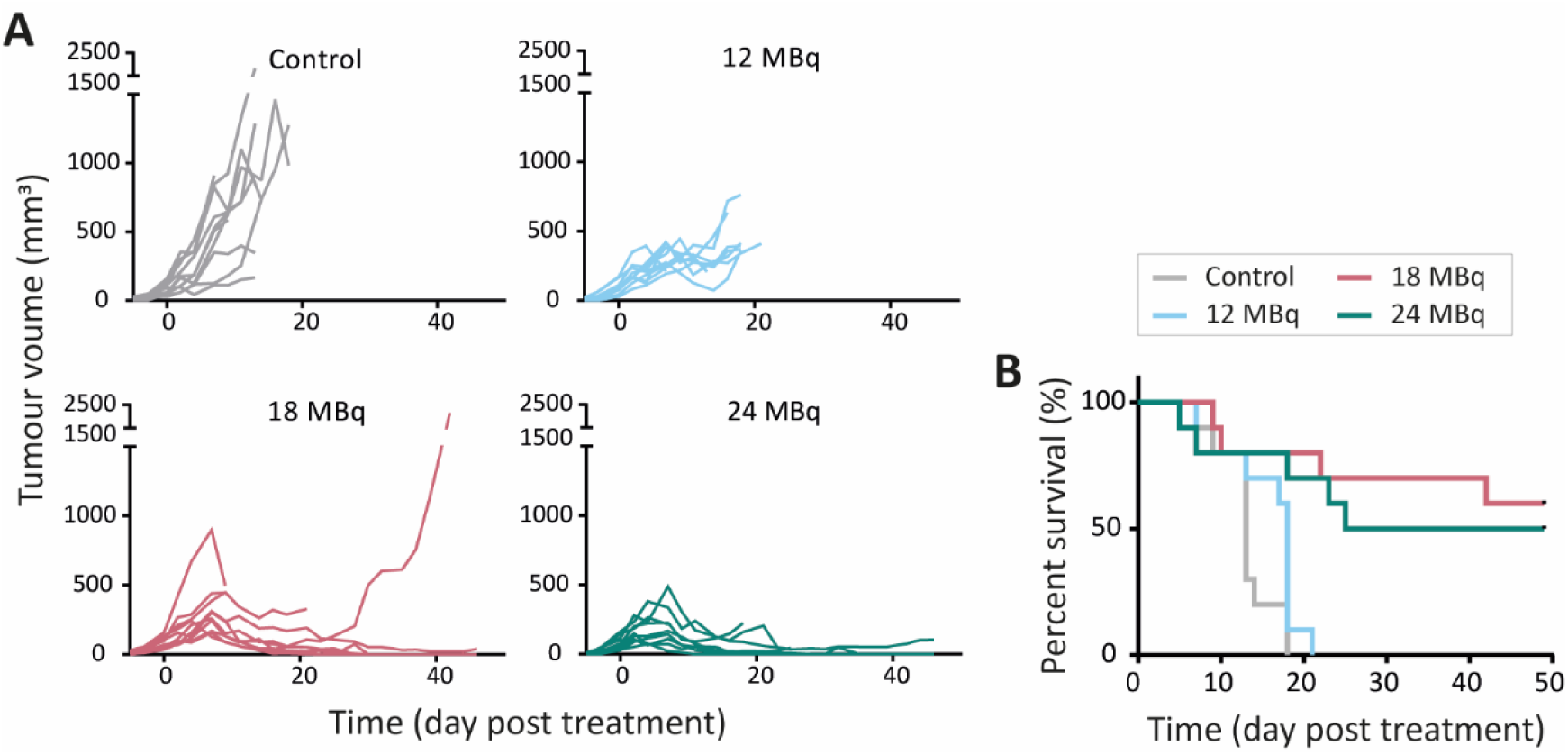
Therapeutic effectiveness of [^177^Lu]Lu-DOTA-hG250 in Renca-CAIX tumor-bearing mice. (**A**) Tumor growth curves of individual mice (n=10/group) after treatment on day 0. (**B**) Survival per group. Significant difference to control group was determined using Kaplan-Meier with log-rank testing and Bonferroni correction for multiple testing (corrected α = 0.0167).

### Complete anti-tumor response upon combined subtherapeutic TRT and ICI

To test the therapeutic efficacy of combined TRT and ICI in poorly T-cell-infiltrated Renca-CAIX tumors, mice received 4 or 12 MBq TRT monotherapy (estimated tumor-absorbed dose of 10 and 29 Gy, respectively), ICI monotherapy (aPD-1 + aCTLA-4), combined TRT/ICI, or vehicle. Combination therapy with 4 or 12 MBq TRT both significantly inhibited tumor growth compared to non-treated animals (p=0.01 and p=0.004), whereas none of the monotherapies showed significant inhibition (**Figure 5A, Table S5**). Survival was significantly improved after 12 MBq TRT (p<0.001) and combination therapy with 4 and 12 MBq TRT (p<0.001) compared to non-treated animals (**Figure 5C**). Also, stable disease and complete tumor responses were most frequently observed after combination therapies (90% and 80% respectively) (**Table S5**). To test for immunological memory, mice with a complete response were re-challenged with Renca-CAIX tumors on one flank and Renca WT tumors on the opposite flank. All mice rejected both Renca-CAIX and Renca WT tumors, except one mouse in the 12 MBq combination therapy group which showed Renca tumor growth (**Table S5**).

**Figure 5.**
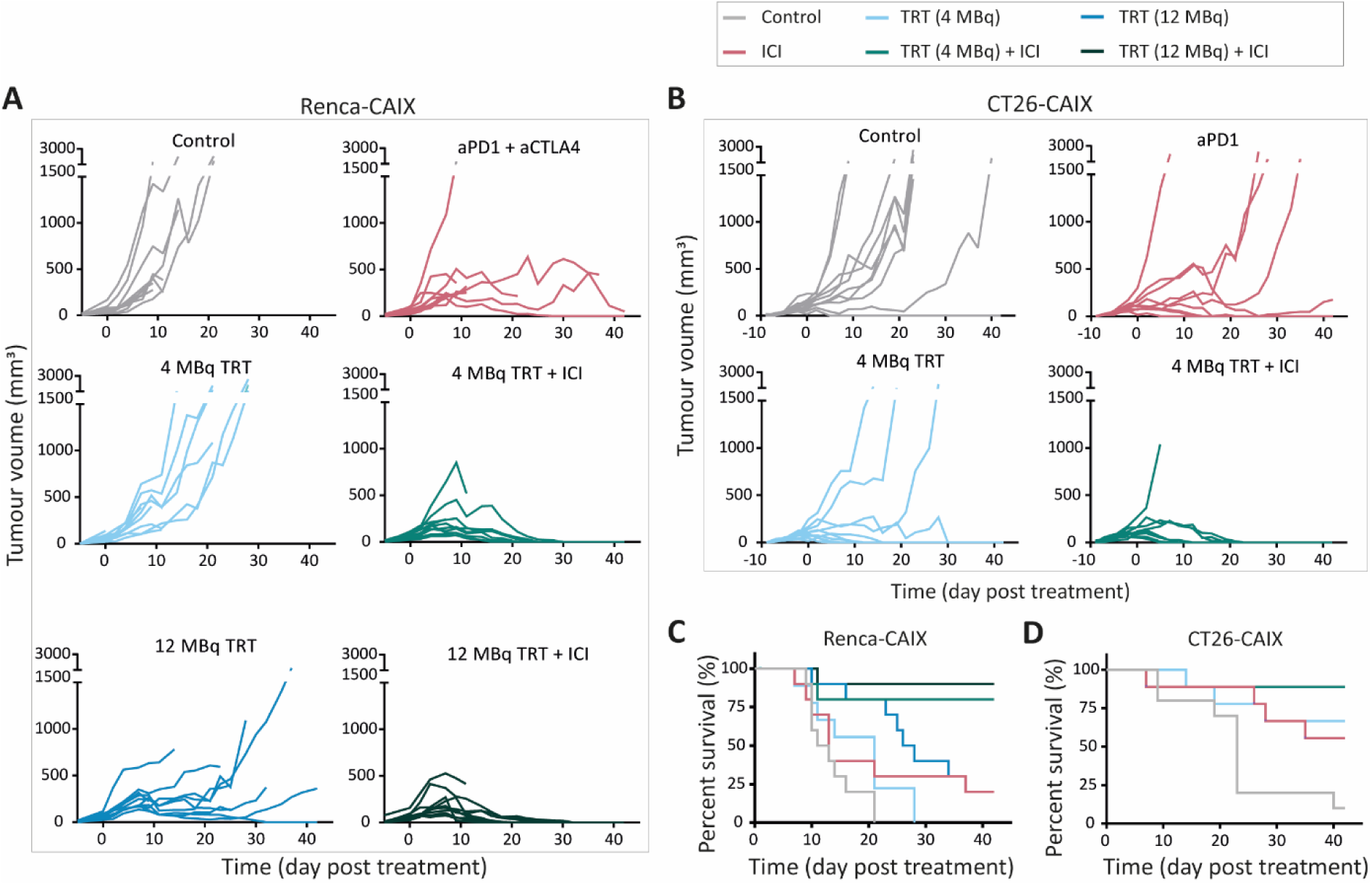
Therapeutic effectiveness of TRT/ICI combination therapy in Renca-CAIX and CT26-CAIX tumor-bearing mice. Tumor growth curves of individual mice after treatment with vehicle (control), aPD-1+aCTLA-4 (ICI), 4 or 12 MBq [^177^Lu]Lu-DOTA-hG250 (TRT), 4 or 12 MBq TRT/ICI on day 0 of Renca-CAIX (n=10/group) (**A**) and treatment with vehicle (control, n=10), aPD-1 (ICI, n=9), 4 MBq TRT (n=9) or 4 MBq TRT/ICI (n=9) on day 0 of CT26-CAIX (**B**) tumor-bearing mice. Survival for Renca-CAIX (**C**) and CT26-CAIX (**D**) tumor-bearing mice.

Based on the former studies, 4 MBq TRT (estimated tumor-absorbed dose of 9 Gy) was selected for future studies and validated in the CT26-CAIX model. However, since these are well-T-cell-infiltrated tumors and generally respond well to ICI treatment [19, 20], TRT was combined with aPD-1 ICI only. In line with the previous experiments, at this dose level only combination therapy resulted in significant tumor growth inhibition (p=0.048) and improved survival (p=0.002) compared to non-treated animals (**Figure 5B**,**D**, **Table S5**). Generally, if a mouse responded to any treatment, this mostly resulted in stable disease or complete tumor regression, and both were most frequently observed after combination therapy (89% and 78% respectively) (**Table S5**). Re-challenge of mice with complete tumor response resulted in rejection of CT26-CAIX tumors in 80-100% of cases, but most mice demonstrated CT26 WT tumor growth (**Table S5**).

Taken together, low dose [^177^Lu]Lu-DOTA-hG250 combined with ICI induced complete responses in approximately 80% of Renca-CAIX and CT26-CAIX tumor-bearing mice.

### TRT/ICI combination therapy alters the Renca-CAIX tumor microenvironment by TRT- and ICI-induced features

To gain insights into the effects of combined [^177^Lu]Lu-DOTA-hG250 and aPD-1/aCTLA-4 therapy on the Renca-CAIX TME, we collected tumor samples on days 0, and 5 and 8 post-treatment. Samples were analyzed through IHC, flow cytometry, and multiplex RNA profiling. Importantly, tumor weights did not significantly differ between the treatment groups at the selected timepoints, so the results observed represented biological changes preceding therapeutic effect (**Figure S6**).

#### Immunohistochemical Analysis

Tumor sections from all treatment groups were (semi)quantitively analyzed for markers of apoptosis (caspase-3), DNA damage (53BP1), cell proliferation (Ki67), CAIX, PD-L1, as well as markers for different immune cell types, including T cells (CD3, CD8, CD4, FOXP3), neutrophils (Ly6G), and macrophages (F4-80) (**Figure 6A-B** and **Figure S7**).

**Figure 6.**
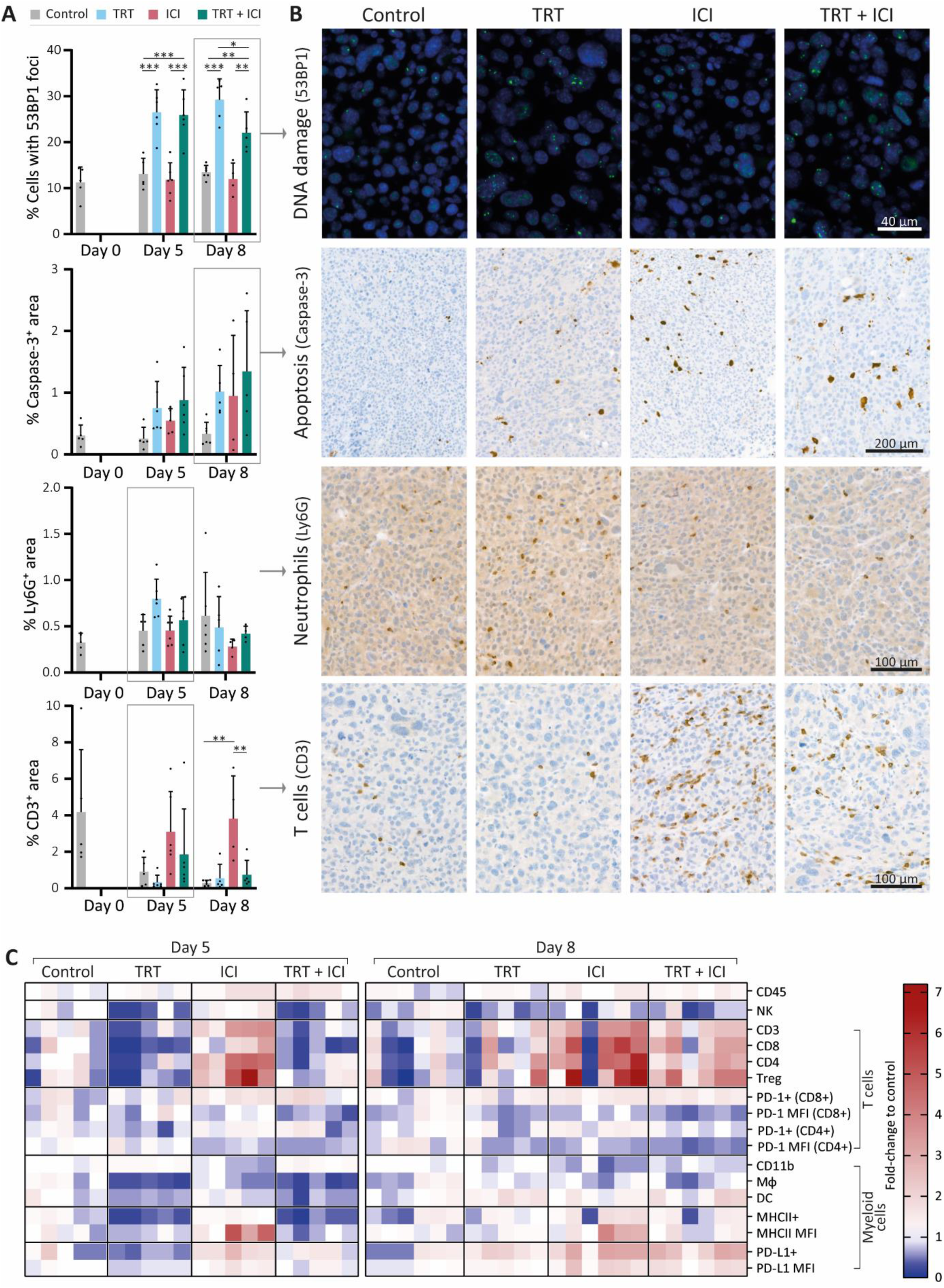
Renca-CAIX tumor-bearing balb/c mice (*n*=5-6/group) were treated with vehicle (control), [^177^Lu]Lu-DOTA-hG250 (TRT), aPD-1 + aCTLA-4 (ICI) or TRT/ICI combination therapy and sacrificed on day 0, 5, or 8 after therapy. (**A**) Quantification of FFPE tumor sections stained for markers of DNA damage (53BP1), apoptosis (caspase-3), neutrophils (Ly6G), or T cells (CD3). Data represents mean + SD. Statistical differences between treatment groups on day 5 and day 8 were separately determined by one-way ANOVA analyses with Šídák’s multiple comparisons test. (*p<0.033, **p<0.01, ***p<0.001). (**B**) Representative images of FFPE tumor sections stained for markers of DNA damage (53BP1, day 8), apoptosis (caspase-3, day 8), neutrophils (Ly6G, day 5), or T cells (CD3, day 5). (**C**) Flow cytometry analyses of tumor-infiltrating immune cells (CD45+), natural killer (NK) cells (CD49b+), T cells (CD3+), cytotoxic T cells (CD3+CD4-CD8+), CD4 T cells (CD3+CD4+CD8-), Tregs (CD3+CD4+CD25+FOXP3+), myeloid cells (CD11b+), macrophages (Mφ, CD11b+F4/80+) and dendritic cells (DC, CD11b+CD11c+) as percent of total immune cells normalized to the mean of control, and expression levels of PD1 (on T cells), MHCII and PD-L1 (on myeloid cells) as mean fluorescent intensity (MFI) normalized to the mean of control. Gating strategies are shown in **Figure S2**.

Generally, Renca-CAIX tumors were increasingly necrotic over time in all groups. Increased apoptosis was observed following each treatment, although not significant, while cell proliferation was significantly increased after TRT and TRT/ICI. Our results most prominently showed that TRT induced DNA damage, as evidenced by a significant increase (p<0.01) in 53BP1^+^ foci-containing cells following both TRT and combination therapy. No apparent differences were observed regarding CAIX or PD-L1 expression or macrophage infiltration among the treatment groups. There was an observable trend in neutrophil counts, albeit not statistically significant, showing an increase following TRT on day 5, supported by pilot studies with higher TRT doses (**Figure S8**). Finally, the strongest evidence was found for T-cell abundance, which was strongly and significantly (p<0.01) increased within the TME following ICI, including all T-cell subtypes (CD8+ cytotoxic, CD4+ T helper, and regulatory T cells). In contrast, TRT caused a minor reduction in T-cell abundance on day 5, while combination therapy resulted in a slight increase on the same day, without reaching statistical significance.

#### Flow cytometry analysis

Flow cytometry analysis enabled quantification of various tumor-infiltrating immune cell subsets relative to the total immune cell population within the tumor. Focusing on the general profile of fold changes compared to control, we observed that the profile of combined TRT/ICI therapy on day 5 was most similar to TRT group, while on day 8 this became more similar to ICI group (**Figure 6C**, **Figure S9**).

We observed an overall increase in the total abundance of immune cells within the tumor after ICI and combination therapy. In the lymphoid compartment, we found a significant decrease of natural killer cells (CD49b^+^) and slight, although non-significant, decreases of various T-cell subsets (CD3^+^, CD8^+^, CD4^+^, Treg (CD4^+^FOXP3^+^CD25^+^)) after TRT and combination therapy on day 5. Conversely, ICI significantly increased the proportion of T cells in the TME. By day 8, the observed decreases following TRT were largely reversed, while ICI continued to enhance T-cell proportions, which was now also observed for combination therapy, although not significantly. The proportion of PD-1^+^CD4^+^ T cells decreased modestly following TRT but increased slightly with ICI, but both not significantly. MFI of PD-1 expression, which may be a proxy for exhaustion, was significantly decreased by both TRT and ICI, although in the case of ICI-treated animals we cannot exclude the possibility that this is partly due to competition with the therapeutically administered anti-PD-1. Within the myeloid compartment, ICI significantly reduced total myeloid (CD11b^+^) cells but left macrophages (CD11b^+^F4-80^+^) and dendritic cells (DCs, CD11b^+^CD11c^+^) unchanged. TRT and combination therapy initially significantly decreased macrophages, DCs, and MHCII^+^ myeloid cells on day 5, but these populations were restored by day 8. Furthermore, the MHCII expression levels increased significantly with ICI, and both PD-L1^+^ myeloid cells and their PD-L1 expression levels were significantly increased on day 8 after ICI and combination therapy. Given the observation of immunological memory in therapeutic studies, we further investigated T-cell recognition by evaluating the proportion of AH1 antigen-specific cytotoxic T cells in both tumor and lymph nodes, as AH1 has been indicated as antigen expressed by Renca tumors [21–23]. In our studies, however, we have found only limited amounts of AH1 antigen-specific cytotoxic T cells, with no significant differences between treatment groups (**Figure S10**).

#### RNA nanostring expression analysis

RNA analysis of tumor sections involved gene expression profiling for 770 genes related to the TME and immune responses using the murine PanCancer IO 360 Gene Expression Panel. Pair-wise comparisons between TRT and control groups and combination and ICI groups, with time as co-variate, identified five gene sets defining the effects of TRT on the TME and five gene sets characterizing the benefits of combination therapy over ICI, based on undirected global significance scores (**Figure S11**).

Gene sets primarily activated by TRT included IFN signaling (e.g. *Ifit1,2,3*, *Irf7*, *Stat1*, *Oas1a*, *Oas3*, *Rsad2*), cytotoxicity, involvement of the lymphoid compartment (e.g. *Cxcl9*, *Cxcl11*, *Gzma*), antigen presentation, and DNA damage repair induction (**Figure 7A,C** and **Figure S12**). Notably, the first four were also prominent in ICI monotherapy and combination therapy, whereas TRT was required for DNA damage repair pathway induction. Within the lymphoid compartment, immune checkpoint genes like *Ctla4, Lag3, Cd274* (PD-L1), and *Icos* were upregulated after ICI and combination therapy, but *Cd86* and *Cd80* and *Icosl* were more elevated following ICI than in combination treatment. Other genes that were more upregulated following ICI treatment compared with combination included *Gata3*, *Prdm1*, and *Ikzf2*. Further comparison highlighted increased cell proliferation and DNA damage repair after combination therapy compared with ICI, aligning with immunohistochemical findings. Conversely, autophagy, MAPK signaling, and metabolic stress signaling pathways were less activated following combination therapy (**Figure 7B-C** and **Figure S12**).

**Figure 7.**
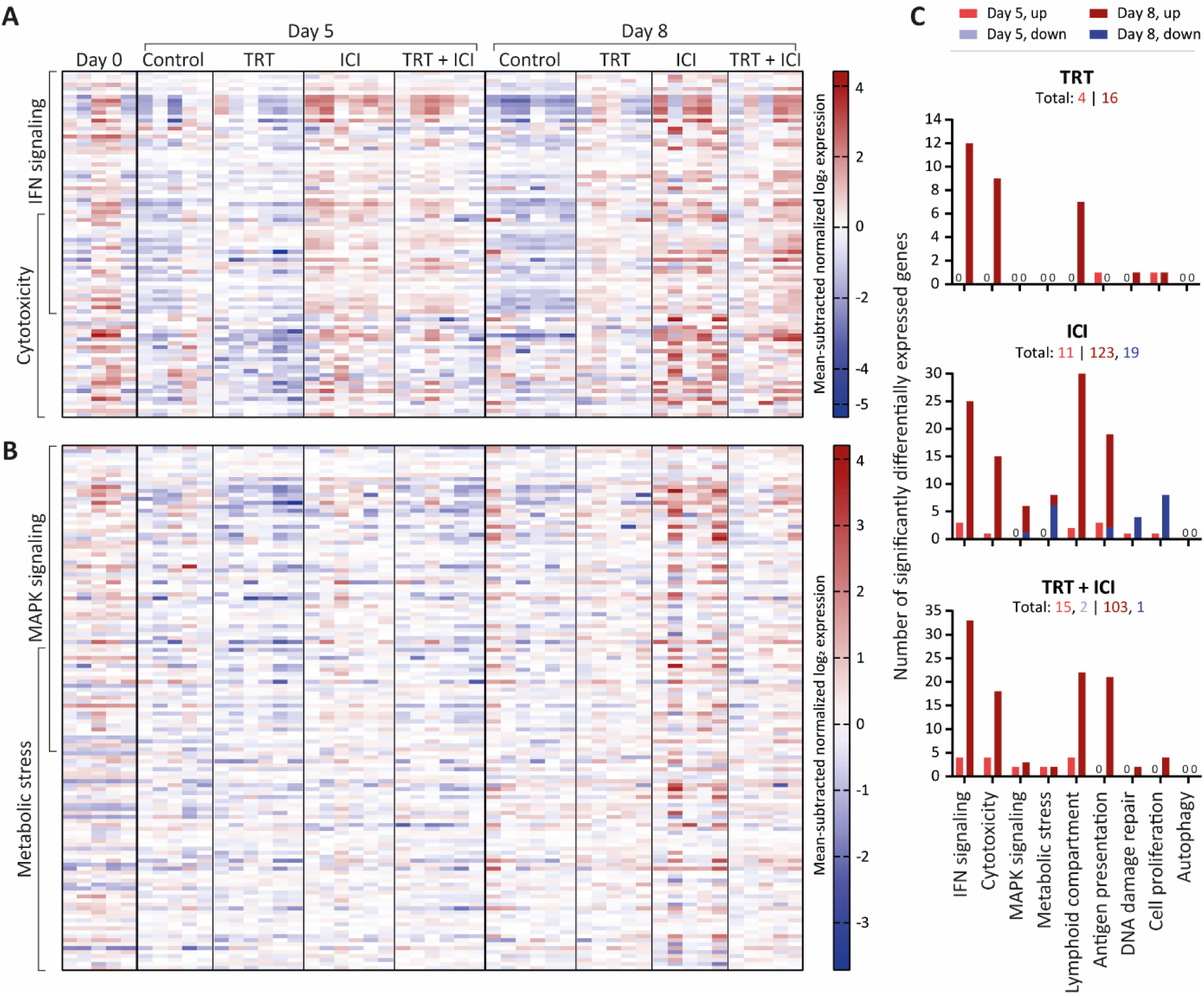
Nanostring gene expression analysis of tumors (n=5-6/group) at 0, 5, or 8 days post-treatment with vehicle (control), [^177^Lu]Lu-DOTA-hG250 (TRT), aPD-1 + aCTLA-4 (ICI) or TRT/ICI combination therapy. (**A**) Heatmap of mean subtracted normalized log2 expression values for all samples of IFN signaling and cytotoxicity gene sets and (**B**) MAPK signaling and metabolic stress gene sets. (**C**) Bar graphs showing numbers of genes significantly upregulated or downregulated for each treatment compared to control groups for the defined gene sets (including those not pictured on the heatmaps in A-B) on day 5 and day 8. Total numbers of up- and downregulated for each group are indicated underneath headings.

Overall, these data reveal both specific and overlapping effects of TRT and ICI on the TME. When TRT and ICI were combined, these effects mostly converged, underscoring the intricate interplay between the two therapies.

## DISCUSSION

Metastatic ccRCC poses a significant challenge, with limited treatment options available and subsequent poor outcome. In this study, we investigated the efficacy and mechanism of action of combined [^177^Lu]Lu-DOTA-hG250 TRT and aPD-1/aCTLA-4 ICI. We utilized two CAIX-expressing murine syngeneic tumor models with comparable intrinsic CAIX expression, radiosensitivity, and *in vivo* [^177^Lu]Lu-DOTA-hG250 biodistribution. However, their PD-L1 expression and T-cell abundance were distinctive, with a higher expected immunogenicity for CT26-CAIX. Combining TRT and ICI induced complete responses in both models and tumor re-challenge studies implied that durable memory immune responses were initiated after treatment.

Exploratory mechanistic studies revealed TRT’s immunomodulatory potential, exemplified by activation of type I IFN signaling and cytotoxicity, while ICI induced even stronger activation of these pathways as well as T-cell tumor infiltration. The TME after combined therapy displayed characteristics of both TRT (e.g. DNA damage induction) and ICI (e.g. T-cell infiltration), while differences with ICI characteristics were observed in RNA expression (e.g. higher cell proliferation, lower PI3K-Akt, and MAPK signaling). These results underscore the rationale of combined TRT/ICI and suggest directions for further exploration, both in terms of efficacy and toxicity.

Favorable efficacy, similar to our findings in combined TRT/ICI, has been reported in previous preclinical studies (recently reviewed in [13, 14]), supporting the proof-of-concept. Although this is cross-experiment comparison, we achieved therapeutic efficacy with [^177^Lu]Lu-DOTA-hG250 TRT monotherapy at a tumor-absorbed dose >29 Gy, whereas similar efficacy could be accomplished with approximately 3-fold lower TRT dose when combined with ICI. Similarly, other studies have shown successful TRT/ICI combination strategies at low tumor-absorbed doses, such as 2.5-5 Gy for ^90^Y [24, 25] and 3.5 Gy for ^177^Lu [26]. In light of these results, future studies will focus on further optimization of dosing levels and frequency. It is important to note that absorbed dose estimations based on biodistribution studies should be interpreted with caution. In our data, we observed considerable variation in tumor uptake of the radiopharmaceutical between different biodistribution and therapy studies, with a trend towards higher tumor uptake in therapy studies (**Figure 3A**, **Figure S13**). This might be explained by radiation-induced changes in the TME upon administration of a therapeutic dose of ^177^Lu. For example, increasing perfusion has been observed even after low-dose irradiation [27, 28], which could affect the tumor uptake and consequently tumor-absorbed doses in therapy studies. Additionally, we observed a slightly lower tumor uptake when [^177^Lu]Lu-DOTA-hG250 was combined with ICI compared with its standalone use (**Figure S13**). Although this effect was not significant, further work with more focus on this difference is warranted.

Furthermore, it is important to examine dose responses in both low and highly immunogenic tumor models. The two models used in this study exhibited comparable intrinsic radiosensitivity, while the *in vivo* response to 9 Gy TRT seemed better for CT26-CAIX. Although this is a cross-experiment comparison, it suggests that biological factors, such as tumor hypoxia, perfusion, or T-cell infiltration, are crucial in predicting TRT and combination therapy responses. Another important factor affecting TRT response was the tumor size at the start of treatment. This was evident in the CT26-CAIX model, where three non-responding mice had the largest tumors (**Figure S14**). For TRT/ICI combination therapy, the response-limiting tumor size was slightly higher, although further comprehensive studies are warranted to evaluate whether larger tumors can be more effectively treated with combination therapy compared to monotherapy.

In contrast to mechanistic insights from other preclinical studies [24, 29–32], the outstanding therapeutic efficacy of combination treatment in our model could not be explained by increased T-cell numbers. However, it must be noted that direct comparison of findings between different studies is challenging due to the time dependency of the observed effects. Still, initial TRT-induced decreases of T cells [24, 30] and absence of prominent T-cell tumor infiltration after combination therapy [33] have been observed previously. Additionally, T-cell numbers do not necessarily reflect T-cell phenotype and functionality, which could be an important question to address in future research. In this context, our RNA profiling did not point to major differences in cytotoxicity, including T-cell cytotoxicity, between combination and ICI treatments. However, flow cytometry demonstrated decreased PD-1 expression on T-cell populations after TRT/ICI, which may indicate less exhausted T cells and corresponds to recent findings in blood samples of patients with progressive metastatic castration-resistant prostate cancer who responded well to combined [^177^Lu]Lu-PSMA-617 and pembrolizumab treatment [34]. Further comparison of our findings with other studies confirms that TRT is a strong inducer of type I IFN signaling [24, 29, 35]. In light of its ability to promote T-cell cross-priming by antigen-presenting cells in context of radiation, there is abundant room for further investigation of the role of IFN in TRT-induced anti-tumor immunity [36]. Along with type I IFN, pre-existing research identified immunogenic cell death hallmarks as TRT-induced immunomodulatory factors, however, in the current study no evidence of increased HMGB1 serum levels was detected (**Figure S15**). Furthermore, our data suggest a role for innate immune cells in TRT/ICI therapy, as we found increased neutrophil tumor infiltration and decreased macrophage populations early after treatment. Further investigations are needed to examine the type of neutrophils and macrophages involved, as antitumor (N1) and protumor (N2) neutrophils have implications in radiation-induced anti-tumor immunity and response to immunotherapy [37–39], and tumor-associated macrophages are known for their tumor-stimulating properties also in ccRCC [40]. Finally, we observed a lowered activity of MAPK signaling following combination therapy compared with ICI, which may have important implications as MAPK inhibition has been associated with ICI response [41]. However, the used methodology was not designed to assess causal relations between therapy and affected pathways, and future studies are therefore recommended. Furthermore, although it is a very useful method for identification of altering pathways, bulk RNA expression profiling gives limited information about specific cell populations and is biased by treatment-induced TME composition alterations. Therefore, single-cell analyses such as single-cell (spatial) transcriptomics may offer a reliable method to further investigate the role of individual cell types.

A limitation of the study is the artificially induced expression of human CAIX in the tumors, which could potentially induce a hCAIX-directed immune response and mask responses to endogenous tumor antigens if hCAIX is immunodominant. To investigate whether the therapy-induced durable memory immune response was directed against hCAIX or not, we included re-challenges with both CAIX+ and CAIX-tumors and confirmed that there was hCAIX-independent immunity in the Renca-CAIX model. Our observation that CT26 WT tumors could develop tumors in the majority of mice, could be a result of the CT26 WT being differently sourced than the CAIX-transfected CT26 WT or alternatively, in this model hCAIX could be the dominant antigen immune responses are directed against. It is worth noting that the development of reliable animal models resembling the TME and immune components observed clinically is generally challenging [42]. Syngeneic murine tumor cell lines come with limitations, including tumor homogeneity and a potential mismatch with the physiological immune environment, particularly in a metastatic setting [43]. The latter is particularly crucial for TRT, due to its promising application in metastatic disease and to the fact that the immune components and mechanisms in the TME of metastasized lesions can differ significantly from primary tumors and each other [42].

Clinically, girentuximab presents a promising radiopharmaceutical for patients with advanced ccRCC given its excellent binding to and specificity for ccRCC lesions [44]. Diagnosis and patient selection can be facilitated accurately by [^89^Zr]Zr-DFO-girentuximab PET imaging [45] and can be used for personalized dosimetry to determine [^177^Lu]Lu-DOTA-girentuximab therapy dosing for individual patients and to study dose-effect relationships in clinical trials. Given the demonstrated therapeutic efficacy of [^177^Lu]Lu-DOTA-girentuximab [10] and the use of ICIs as a standard of care treatment, the implementation of the combined treatment as clinical treatment for advanced ccRCC is within reach, which is underscored by two ongoing clinical trials (NCT05663710, NCT05239533) [46, 47], of which the results are eagerly awaited. Particularly the opportunity for dose-reduction can contribute to a better treatment, including lower myelotoxicity of TRT which is supported by observations for the lowest administered dose of [^177^Lu]Lu-DOTA-girentuximab in a phase I study [11]. For ICI, dose-reduction may minimize immune-related adverse events commonly associated with the treatment.

This investigation aimed to gain a proof-of-concept of combined [^177^Lu]Lu-DOTA-hG250 and aPD-1/aCTLA-4 therapy using syngeneic mouse models and to explore its radiobiological and immunological effects on the TME. Our results demonstrate the outstanding therapeutic efficacy of the combination treatment using subtherapeutic [^177^Lu]Lu-DOTA-hG250, including complete remissions and durable memory immune responses. Furthermore, these studies show that the combination treatment is accompanied by TRT-induced DNA damage, cell damage, cell death, and type I IFN signaling and ICI-induced immune activation exemplified by T-cell infiltration and cytotoxicity. Overall, this study supports the use of combined TRT/ICI for patients with advanced ccRCC. Furthermore, our findings shed new light on the biological effects of TRT, ICI, and combination treatment on the TME. This provides directions for further mechanistic preclinical studies, which can contribute to our understanding of combined TRT/ICI treatment and thus assist the design of future clinical trials and clinical implementation of the treatment to realize its full potential.

## DECLARATIONS

## Acknowledgements

We thank Sylvia Wenker, Sanne van Lith, and Peter Laverman for valuable discussions. We thank Floor Moonen, Bianca Lemmers-van de Weem, Karin de Haas-Cremers, Kitty Lemmens-Hermans, and Liz van den Brand for technical assistance with the animal experiments, and Gerwin Sandker for co-creating the IHC analysis macro. (Parts of) this study has been presented at the KWF-Dutch Tumor Immunology Meeting 2023, the European Association of Nuclear Medicine 2022 and 2023 conferences, the European Association of Urology 2023 conference, and the 2^nd^ workshop on Radiobiology of Molecular Radiotherapy 2023 conference and published in the Abstract Books.

## Author’s contributions

Conceptualization: SCK, EO, MWK, MPW, KTB, SH. Data collection: SCK, JM, CF, GF, DB, MG, MH, DK, JH. Data analysis: SCK, GT, MWK, KTB. Writing – original draft preparation: SCK. Writing – review and editing: all authors.

## Ethics approval

All *in vivo* experiments were approved by the Animal Welfare Body of the Radboud University, Nijmegen, and the Central Authority for Scientific Procedures on Animals (AVD1030020209645) and were performed in accordance with the principles stated by the Dutch Act on Animal Experiments (2014).

## Patient consent for publication

Not applicable

## Data availability statement

The data generated and/or analyzed during this study are available from the corresponding author upon reasonable request.

### Abbreviations

%IA/g: percentage injected activity per gram
BSA: Bovine Serum Albumin
CAIX: Carbonic Anhydrase IX
ccRCC: Clear Cell Renal Cell Carcinoma
CTLA-4: Cytotoxic T Lymphocyte Antigen-4
DOTA: S-2-(4-Isothiocyanatobenzyl)-1,4,7,10-tetraazacyclododecane tetraacetic acid (p-SCN-Bn-DOTA)
EBRT: External Beam Radiation Therapy
EDTA: ethylenediaminetetraacetic
FFPE: Formalin-fixed paraffin-embedded
hG250: humanized monoclonal antibody G250 (girentuximab)
HMGB1: High-Mobility Group Box 1
ICI: Immune Checkpoint Inhibitors
IFN: Interferon
IHC: Immunohistochemistry
MBq: Megabequerel
MHC: Major Histocompatibility Complex
nAUC: normalized Area Under the Curve
PBS: phosphate buffered saline
PD-1: Programmed Cell Death Receptor-1
PD-L1: Programmed Cell Death Receptor-1 Ligand
RT: room temperature
TME: Tumor Microenvironment
TRT: Targeted Radionuclide Therapy
VHL: Von Hippel Lindau

