## Supplemental Material for "Towards Effective CAIX-targeted Radionuclide and Checkpoint Inhibition Combination Therapy for Advanced Clear Cell Renal Cell Carcinoma"

### SUPPLEMENTAL METHODS

#### Conjugation, radiolabeling, and quality control

hG250 was conjugated with S-2-(4-Isothiocyanatobenzyl)-1,4,7,10-tetraazacyclododecane tetraacetic acid (p-SCN-Bn-DOTA)(Macrocyclics™, Plano) by adding DOTA to hG250 at a 30:1 mass ratio in 0.1 M $NaHCO_3$ pH 9.5 for 1h at RT, and subsequently dialyzed against 0.25 M $NH_4Ac$ pH 5.5 supplemented with Chelex resin (2 g/L, Bio-Rad Laboratories) [1]. [$^{111}$In]InCl$_3$ was obtained from Curium (Petten, The Netherlands) and no-carrier-added [$^{177}$Lu]LuCl$_3$ was obtained from ITM Medical Isotopes GmbH (Garching, Germany). DOTA-conjugated hG250 was incubated with [$^{111}$In]InCl$_3$ or [$^{177}$Lu]LuCl$_3$ in 0.5 M MES buffer, pH 5.5 under metal-free conditions at 37°C for 30 min at 5 MBq/µg unless indicated otherwise. After incubation, labelling efficiency was determined by instant layer chromatography (ITLC), using ITLC silica gel strip (Agilent Technologies) and 0.1 M citrate buffer pH 6.0, and 50 mM ethylenediaminetetraacetic acid (EDTA) was added. If labelling efficiency was <90% for *in vitro* experiments and <95% for *in vivo* experiments, the reaction mixture was purified using PD-10 desalting columns (VWR, 17-0851-01) eluted with 0.9% NaCl. For animal studies, unlabeled hG250 was added to the injection solution to achieve the desired protein concentration. The immunoreactive fraction, determined as described by Lindmo et al., exceeded 70% for all preparations [2].

##### *In* vitro flow cytometry

CAIX antigen expression of all cell lines was checked using flow cytometry. Cells were stained with hG250 (1 ug/ml, 30 min, 4°C) in PBA (PBS, 0.5% BSA, 0.05% sodium azide) and goat-anti-human AlexaFluor488 labelled antibody (ThermoScientific, A-11013, 1:300, 15 min, 4°C). Cells were analyzed on a FACSCanto II Flow Cytometry System (BD Biosciences) and results were analyzed with FlowJo 10.8.2 (FlowJo LCC).

#### Cell viability assay

Radiosensitivity to TRT was assessed using cell viability assays using the CellTiter-Glo assay (Promega, G7570) according to manufacturer's instructions. Cells were seeded in 96-well plates to attach overnight, treated for 24h with 0-6 MBq/mL [$^{177}$Lu]Lu-DOTA-hG250 in 200 µL culture medium, and cultured for another 8 days in fresh medium. Luminescence values were normalized to untreated cells and the data was described using the log(inhibitor)-response variable slope model with constrained bottom (0.0) and top (1.0) values. Differences in $IC_{50}$ between cell lines was statistically tested using one-way ANOVA with Tukey's multiple comparison correction.

#### Animal experiments

Protocols for each experiment were registered at preclinicaltrials.eu (PCTE0000440- PCTE0000445). For each animal experiment, the number of animals injected with tumor cells was determined accounting for an expected tumor take rate of 83% for Renca-CAIX and 71% for CT26-CAIX. For *ex vivo* biodistribution studies, sample size (*n*=5/group, 30 total) was selected based on previous biodistribution studies [3], but there was a dropout of 8 animals due to low tumor take resulting in the group sizes presented in **Table S3**. A priori sample size calculations for therapy studies were based on an effect size of 75% decline in normalized area under the curve (nAUC) compared to control, significance level of 5% and power of 80%. This resulted in a sample size of 10 mice per group for all therapy experiments. In the experiment with CT26-CAIX tumor-bearing mice, failed tumor cell injection causing intraperitoneal tumor growth resulted in 3 dropouts, resulting in the group sizes presented in **Table S4**. Experiments for *ex vivo* analyses were determined on 5 (endpoint day 0 or 5) or 6 (endpoint day 8, to account for possible humane endpoints). Initially, nine extra mice were included, because

tumor take was higher and humane endpoints were lower than expected. However, two mice were excluded from analyses because of failed tumor injection and complete tumor remission, resulting in the group sizes presented in the heatmaps of **Figures 6-7**.
Selection of mice for each groups was determined by block randomization based on tumor volume, using a random number generator. Mice from different groups were housed together to minimize confounding effects and cages were stored randomly. The order in which mice were treated and measured was random, as well as the measurements of tumor microenvironment characteristics ((immunohistochemistry, flow cytometry, RNA analysis).
Biotechnicians, who performed all injections and measurements and determined whether an animal must be sacrificed due to a humane endpoint, were blinded for group allocation. The investigator was blinded for group allocation during assessment of the tumor microenvironment (immunohistochemistry, flow cytometry, RNA analysis), by concealing of animal number.

**Dosimetry estimations**

Time-activity curves (TAC) for tumors were fitted to a power law function followed by a mono-exponential decay in Python. Tumor growth was modelled with a mono exponential function and used to correct both bound activity (%IA/g) over time and define a time-dependent S-value (e.g. absorbed dose rate per unit activity), evaluated by Geant4 11.02 as previously reported [4]. The tumor-absorbed dose was calculated by numerical integration of the dose-rate curve (**Figure S1**). The absorbed dose rate error was determined by propagating uncertainties from activity measurements and tumor growth curve fitting, affecting both bound activities and S-value. The difference between the areas under the dose rate error bounds was used to calculate the uncertainty in tumor-absorbed doses. The time-integrated activity coefficient for livers was calculated according to the trapezoid integration method [5]. Normal organ S-values for $^{177}$Lu were obtained from OLINDA/EXM 2.0 and multiplied with obtained TIACs for each organ to calculate absorbed doses according to Medical Internal Radiation Dose (MIRD) Committee methodology [6, 7].

**Pilot animal experiment neutrophil quantification**

Biological effects of [$^{177}$Lu]Lu-DOTA-hG250 at different doses was studied in Renca-CAIX tumor-bearing mice (*n*=3/group, 12 total). Mice were sacrificed at 7 days post treatment with 5 µg of 12, 18, or 24 MBq [$^{177}$Lu]Lu-DOTA-hG250 or vehicle control (0.9% NaCl) and tumors were harvested, fixed in 4% formalin and embedded in paraffin (FFPE).

***Ex vivo* flow cytometry**

For panel 3, cells were first stained with AH1 dextramer according to the manufacturer's instructions. For all panels, cells were incubated with live/death marker in PBS for 20 min on ice, and subsequently incubated for 10 min on ice with CD16/CD32 (1:800, BD, 553142) for Fc blocking. Cells were incubated with the given extracellular antibodies (**Table S2**) for 30 min on ice. For panel 1, cells were incubated with BV510-streptavidin secondary antibody (1:300, BD, 563261) for 15 min on ice. For all panels, cells were fixed for 30 min on ice (foxp3/transcription factor staining buffer set, Invitrogen, 00-5523). For panel 2, cells were incubated with FoxP3 antibody for 30 min at RT. Samples were analyzed on a FACSCanto II Flow Cytometry System (BD Biosciences) and results were analyzed with FlowJo using the specified gating strategies (**Figure S2**).

**Immunohistochemistry and quantification**

FFPE tumor sections were deparaffinized and rehydrated, followed by antigen retrieval in 10 mM sodium citrate pH 6.0 (caspase-3, Ki67, CAIX, CD4) or TBE (VWR) + 0.05% Tween-20 (53BP1, FOXP3) using a PT module (ThermoScientific) with a 10 min 96°C protocol. For 53BP1, sections were permeabilized with 0.5% Triton-X-100 for 20 min and blocked by 1h incubation in 0.1% Triton-X-100+2% bovine serum albumin (BSA). For the other stainings, endogenous peroxidase activity was blocked by 10 min incubation in 3% $H_2O_2$ and non-specific Ig binding was blocked by 30 min incubation in 20% normal goat serum (5095, Bodinco) or normal rabbit serum (FOXP3 (5096, Bodinco)). Sections were incubated with primary antibodies at RT (diluted in 1% BSA in PBS or 0.1% Triton+2% BSA in PBS (53BP1)): rabbit-anti-53BP1 (1:500, 1.5h, Novus biologicals, NB100-904), rabbit-anti-caspase-3 (1:4000, O/N at 4°C, Vector, BA-1000), rabbit-anti-Ki67 (1:150, 1h, ThermoScientific, RM-9106-S1), rabbit-anti-CAIX (1:1000, 1h, Novus Biologicals, NB100-417), rabbit-anti-CD4 (1:2000, 1h, Abcam, ab183685), rat-anti-FOXP3 (1:400, 1h, ThermoScientific, 14-5773-82). Slides were incubated with secondary antibody at RT: goat-anti-rabbit-Alexa488 (1:500, 1h at 4°C, Invitrogen, A-11008), goat anti-rabbit-biotin (1:200, 30 min, Vector Laboratories, BA-1000), rabbit-anti-rat-biotin (1:200, 30 min, Vector Laboratories, BA-4001). For biotinylated secondary antibodies, slides were incubated with Vectastain ABC-HRP kit according to manufacturer’s instructions (Vector Laboratories, PK-6100), before 8 min incubation with Bright DAB solution (Immunologic, B-500), counterstaining with hematoxylin, dehydration, and mounting with a cover slip using permount (ThermoScientific, 15832544). For 53BP1, sections were counterstained with DAPI (1:2000, Life technologies, D1306) and mounted with a cover slip using fluoromount (Sigma, F4680). Necrotic area was estimated from H&E-stained tumors manually. Intratumoral immune cell presence was quantified using; color deconvolution (HDAB), manual threshold (6,255) and pixel count for hematoxylin to determine the total analyzed area, and auto threshold “Yen” and analyze particles (size=10-∞) for DAB. $DAB^+$ and total areas were summed for all analyzed images of one tumor, $DAB^+$ areas were divided by total areas to obtain a percentage of positive area for each tumor. For Ki67, DAB+ and hematoxylin area were determined using auto threshold “Otsu” and analyze particles (size=10-400, and size=10-∞, respectively) to obtain a ratio between $DAB^+$ and $hematoxylin^+$ areas. For fluorescent signals, thresholds for segmentation were manually set above background, after which 53BP1 and DAPI signal was quantified from binary images using particle analysis.

#### SUPPLEMENTAL TABLES

**Table S1.** Details on used cell lines.

| Cell Line | Type | Supplier |
|---|---|---|
| SKRC-52 | Human renal-cell carcinoma | Memorial Sloan Kettering Cancer Center, CVCL_6198 |
| CT26 | Murine colorectal carcinoma | ATCC, CRL-2638 |
| CT26-CAIX | hCAIX-transfected murine colorectal carcinoma | Kindly provided by Professor D. Neri (Swiss Federal Institute of Technology, Zürich, Switzerland [8]) |
| Renca | Murine renal carcinoma | ATCC, CRL-2947 |
| Renca-CAIX | hCAIX-transfected murine renal carcinoma | In house. Transfection of Renca cells by cloning hCAIX cDNA into the pBJ1-Neo mammalian expression vector (Addgene:1923) containing G418, using FuGENE6 transfection reagent (FuGENE) according to manufacturer's instructions. Selection of positive clones with base medium supplemented with G418. |

**Table S2.** Antibody panels for flow cytometry

| **1: myeloid panel** | | | | |
|---|---|---|---|---|
| **Marker** | **Fluorophore** | **Dilution** | **Catalog nr.** | **Company** |
| Fixable viability dye | eFluor 450 | 1:4000 | 65-0863-18 | eBioscience |
| CD11b-biotin | | 1:200 | 13-0112-85 | eBioscience |
| PD-L1 | Alexa Fluor 488 | 1:100 | 53-5982-82 | Invitrogen |
| MHCII | PE | 1:800 | 1138040 | Antibodychain |
| CD45.2 | PerCPCy5.5 | 1:200 | 552950 | BD |
| F4/80 | PeCy7 | 1:150 | 1215570 | Antibodychain |
| CD11c | APC | 1:400 | 1186550 | Antibodychain |

| **2: T cell panel** | | | | |
|---|---|---|---|---|
| **Marker** | **Fluorophore** | **Dilution** | **Catalog nr.** | **Company** |
| Fixable viability dye | eFluor 450 | 1:4000 | 65-0863-18 | eBioscience |
| CD3e | BV510 | 1:50 | 100353 | Biolegend |
| CD8β.2 | FITC | 1:1000 | 1302020 | Antibodychain |
| PD-1 | PE | 1:50 | 12-9985-83 | eBioscience |
| CD45.2 | PerCPCy5.5 | 1:200 | 552950 | BD |
| FoxP3 | PeCy7 | 1:200 | 25-5773-82 | eBioscience |
| CD25 | APC | 1:400 | 17-0251-82 | eBioscience |
| CD4 | APCCy7 | 1:200 | 1102630 | Antibodychain |

| **3: AH antigen-specific T cell panel** | | | | |
|---|---|---|---|---|
| **Marker** | **Fluorophore** | **Dilution** | **Catalog nr.** | **Company** |
| Fixable viability dye | eFluor 450 | 1:4000 | 65-0863-18 | eBioscience |
| CD3e | BV510 | 1:50 | 100353 | Biolegend |
| CD8b.2 | FITC | 1:1000 | 1302020 | Antibodychain |
| CD45.2 | PerCPCy5.5 | 1:200 | 552950 | BD |
| AH1 dextramer | APC | | JG03294-APC | Immudex |
| CD49b | APCCy7 | 1:50 | A15420 | Invitrogen |

**Table S3.** Statistical parameters for radiosensitivity assays. α/β ratios derived from a linear quadratic model non-linear regression fit to clonogenic survival data and surviving fraction at 2 Gy (SF2) after EBRT and IC50 values after TRT are shown. Data represents mean ± SEM of 2 independent experiments and p-values are given for comparison between the cell lines using one-way ANOVA.

| | **SKRC-52** | **Renca-CAIX** | **CT26-CAIX** |
|---|---|---|---|
| **EBRT clonogenic survival – α/β ratio** | 9.02 ± 3.84 | 3.03 ± 1.56 | 1.86 ± 0.75 |
| Compared to control | | p=0.21 | p=0.12 |
| Compared to CT26-CAIX | | p=0.94 | |
| **EBRT clonogenic survival – SF2** | 0.35 ± 0.04 | 0.61 ± 0.07 | 0.65 ± 0.05 |
| Compared to control | | p=0.08 | p=0.06 |
| Compared to CT26-CAIX | | p=0.85 | |
| **TRT cell viability – IC50** | -0.05 ± 0.05 | 0.71 ± 0.09 | 0.58± 0.14 |
| Compared to control | | p<0.001 | p<0.001 |
| Compared to CT26-CAIX | | p=0.63 | |

**Table S4.** *Ex vivo* biodistribution analysis of [$^{177}$Lu]Lu-DOTA-hG250 in Renca-CAIX and CT26-CAIX tumor-bearing BALB/cAnNRj mice at 1, 3, and 7 days post injection. Data represent mean tissue uptake (in %IA/g) with standard deviation.

| **Renca-CAIX** | **1 day.** (n=5) | **3 days** (n=5) | **7 days** (n=4) |
|---|---|---|---|
| | (%IA/g) | (%IA/g) | (%IA/g) |
| **Blood** | 12.1 ± 2.7 | 4.3 ± 1.4 | 1.5 ± 1.0 |
| **Heart** | 3.6 ± 1.2 | 1.4 ± 0.4 | 0.6 ± 0.3 |
| **Lung** | 9.8 ± 3.9 | 4.0 ± 1.4 | 1.3 ± 0.6 |
| **Liver** | 33.8 ± 8.8 | 38.9 ± 7.9 | 20.3 ± 1.4 |
| **Spleen** | 16.8 ± 5.2 | 10.5 ± 2.5 | 5.4 ± 0.9 |
| **Pancreas** | 1.8 ± 0.5 | 0.9 ± 0.1 | 0.5 ± 0.2 |
| **Kidney** | 4.8 ± 1.4 | 3.1 ± 0.4 | 1.8 ± 0.2 |
| **Stomach** | 1.9 ± 0.4 | 0.8 ± 0.1 | 0.5 ± 0.2 |
| **Small intestine** | 4.2 ± 1.3 | 2.1 ± 0.6 | 0.9 ± 0.3 |
| **Muscle** | 0.8 ± 0.2 | 0.4 ± 0.1 | 0.3 ± 0.1 |
| **Bone** | 3.0 ± 1.1 | 1.5 ± 0.5 | 1.1 ± 0.4 |
| **Tumor** | 31.6 ± 9.4 | 22.0 ± 9.4 | 9.7 ± 3.2 |

| **CT26-CAIX** | **1 day** (n=3) | **3 days** (n=2) | **7 days** (n=3) |
|---|---|---|---|
| | (%IA/g) | (%IA/g) | (%IA/g) |
| **Blood** | 10.3 ± 1.9 | 5.1 ± 1.4 | 3.2 ± 1.1 |
| **Heart** | 2.8 ± 0.3 | 1.4 ± 0.3 | 1.0 ± 0.2 |
| **Lung** | 7.3 ± 1.5 | 4.0 ± 0.7 | 2.6 ± 0.8 |
| **Liver** | 34.7 ± 6.5 | 27.2 ± 4.0 | 14.9 ± 1.5 |
| **Spleen** | 15.0 ± 1.1 | 8.7 ± 0.0 | 6.5 ± 0.3 |
| **Pancreas** | 1.4 ± 0.2 | 1.0 ± 0.4 | 0.6 ± 0.1 |
| **Kidney** | 4.2 ± 0.2 | 2.8 ± 0.4 | 1.9 ± 0.2 |
| **Stomach** | 1.3 ± 0.2 | 0.9 ± 0.2 | 0.6 ± 0.1 |
| **Small intestine** | 3.0 ± 1.1 | 2.2 ± 0.1 | 1.3 ± 0.2 |
| **Muscle** | 0.6 ± 0.1 | 0.5 ± 0.0 | 0.3 ± 0.1 |
| **Bone** | 1.9 ± 0.6 | 1.4 ± 0.2 | 2.1 ± 0.4 |
| **Tumor** | 16.8 ± 2.3 | 16.1 ± 6.0 | 14.0 ± 3.7 |

**Table S5.** Results of three therapy experiments using [$^{177}$Lu]Lu-DOTA-hG250 TRT with or without ICI in Renca-CAIX or CT26-CAIX tumor-bearing mice. Sample size per group, median normalized area under the curve (nAUC) with interquartile range (IQR), adjusted P-values after pair-wise comparisons using Kruskal-Wallis test with Dunn's multiple comparison correction test, and percentages of animals with stabilized disease, complete response, and tumor rejection of CAIX+ and CAIX- tumors after re-challenge are reported.

| **Group** (sample size) | **nAUC** ("Median (IQR)" ($mm^3$)) | **Adjusted P-value nAUC** (Kruskal-Wallis test with Dunn's multiple comparison) | **P-value survival** (Mantel-Cox log-rank testing with Bonferroni correction) | **Stable disease** (%) | **Complete response** (%) | **CAIX+ tumor rejection** (%) | **CAIX- tumor rejection** (%) |
|---|---|---|---|---|---|---|---|
| **Therapy dosing study Renca-CAIX** | | | | | | | |
| Control (n=10) | "329(225)" | | Adjusted α=0.0167 | 0 | 0 | | |
| 12 MBq (n=10) | "201(51)" | 0.64 | 0.09 | 0 | 0 | | |
| 18 MBq (n=10) | "89(204)" | 0.03 | 0.001 | 70 | 50 | | |
| 24 MBq (n=10) | "61(101)" | 0.0002 | 0.004 | 50 | 40 | | |
| **Combination therapy study Renca-CAIX** | | | | | | | |
| Control (n=10) | "270(320)" | | Adjusted α=0.0056 | 0 | 0 | | |
| 4 MBq TRT (n=10) | "414(280)" | >0.99 | 0.10 | 11 | 0 | | |
| 12 MBq TRT (n=10) | "166(179)" | >0.99 | 0.0003 | 40 | 10 | 100 | 100 |
| ICI (n=10) | "161(134)" | >0.99 | 0.22 | 20 | 10 | 100 | 100 |
| 4 MBq TRT + ICI (n=10) | "58(70)" | 0.01 (vs control)<br>0.006 (vs TRT)<br>0.23 (vs ICI) | 0.0009 (vs control)<br>0.002 (vs TRT)<br>0.018 (vs ICI) | 80 | 80 | 100 | 100 |
| 12 MBq TRT + ICI (n=10) | "55(61)" | 0.004 (vs control)<br>0.046 (vs TRT)<br>0.08 (vs ICI) | 0.0001 (vs control)<br>0.003 (vs TRT)<br>0.003 (vs ICI) | 100 | 80 | 100 | 88 |
| **Combination therapy study CT26-CAIX** | | | | | | | |
| Control (n=10) | "344(161)" | | Adjusted α=0.01 | 10 | 10 | 100 | 0 |
| 4 MBq TRT (n=9) | "26(313)" | 0.12 | 0.02 | 89 | 67 | 80 | 0 |
| aPD1 (n=9) | "173(351)" | >0.99 | 0.03 | 44 | 44 | 100 | 50 |
| TRT + ICI (n=9) | "23(42)" | 0.048 (vs control)<br>>0.99 (vs TRT)<br>0.64 (vs ICI) | 0.002 (vs control)<br>0.31 (vs TRT)<br>0.15 (vs ICI) | 89 | 78 | 100 | 17 |

#### SUPPLEMENTAL FIGURES

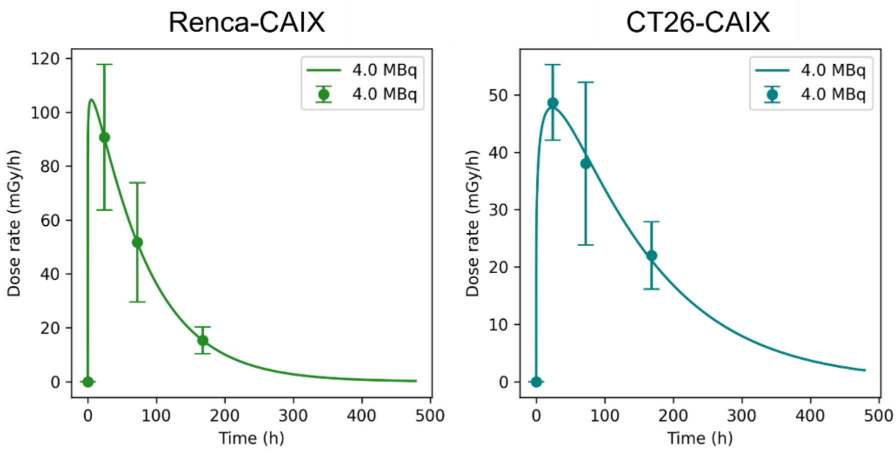


**Figure S1.** Dose-rate curves for Renca-CAIX (left panel) and CT26-CAIX (right panel) tumors, which were integrated to obtain tumor-absorbed doses.

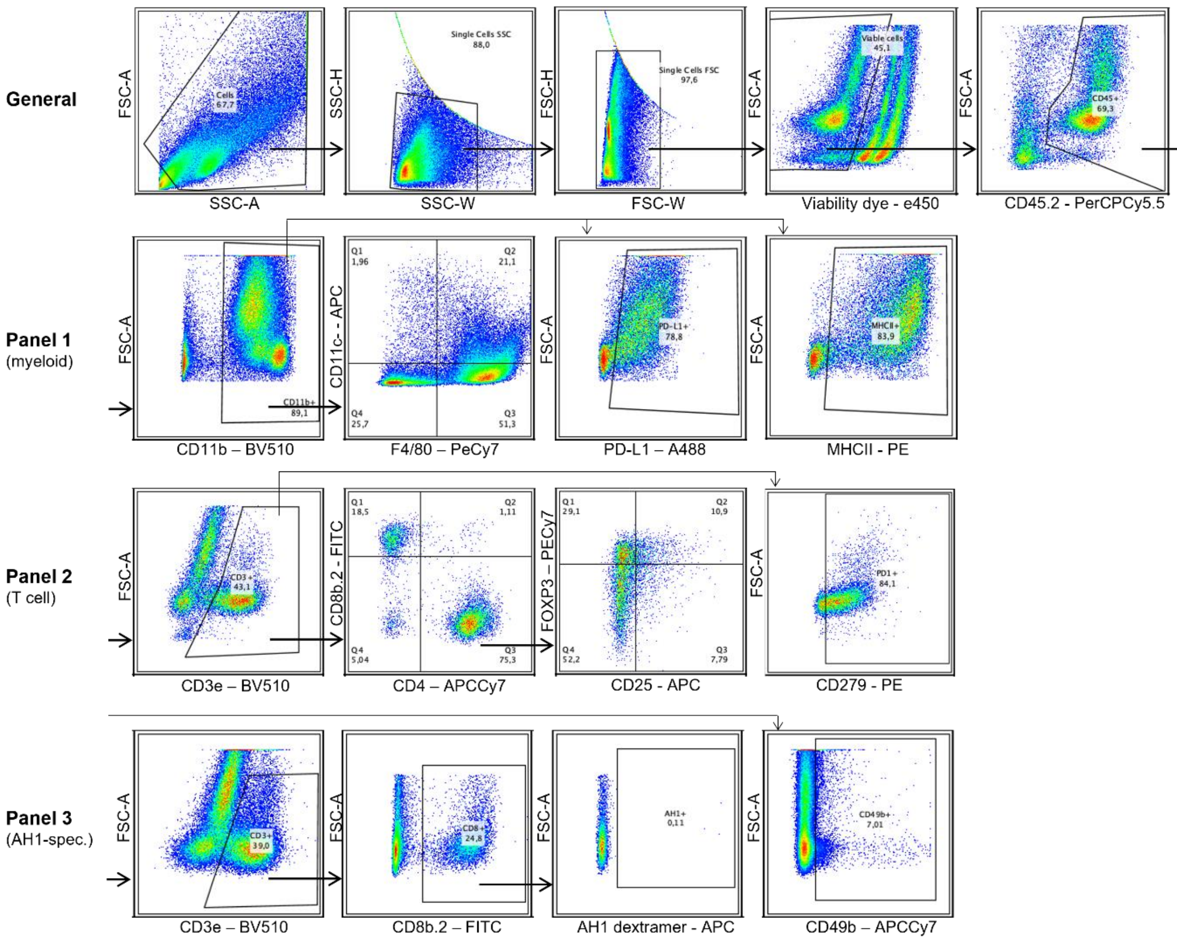


**Figure S2.** Gating strategies for all flow cytometry panels. Gating for all panels started with general gating (upper panels) followed by panel-specific gating as indicated. Gate setting for each marker was based on Fluorescence Minus One (FMO) controls

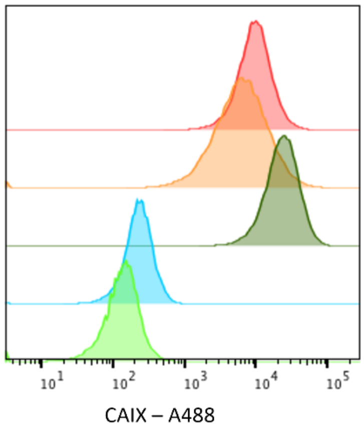


| Cell Line | Mean Fluorescent Intensity: CAIX |
|---|---|
| Renca-CAIX | 8773 |
| CT26-CAIX | 5597 |
| SKRC-52 | 21150 |
| Renca WT | 218 |
| CT26 WT | 122 |

**Figure S3**. CAIX expression on all used cell lines, evaluated by flow cytometry. CAIX expression is presented as histogram and mean fluorescent intensity value.

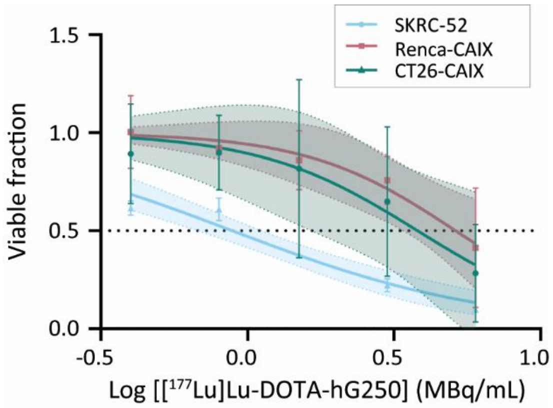


**Figure S4**. In vitro radiosensitivity to TRT. Viability of cells after 24 h of treatment with 0-6 MBq/mL [$^{177}$Lu]Lu-DOTA-hG250 and 8 days additional culturing. Data represent mean ± SD of 2 independent experiments. Non-linear regression using the log(inhibitor)-response variable slope model was used to fit the data and 95% confidence bands are shown (dashed lines).

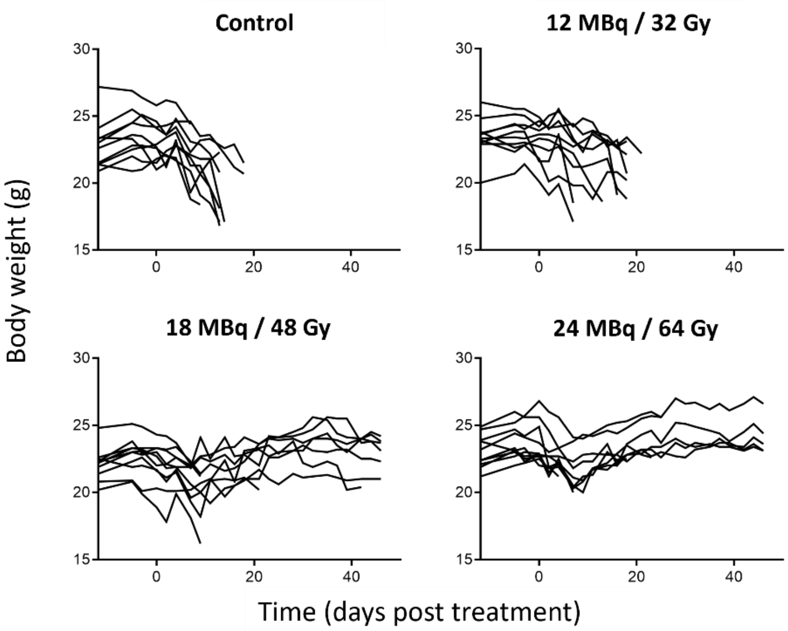


**Figure S5**. Body weight curves of individual mice after treatment with vehicle (control), 12, 18, or 24 MBq [$^{177}$lu]Lu-DOTA-hG250 on day 0.

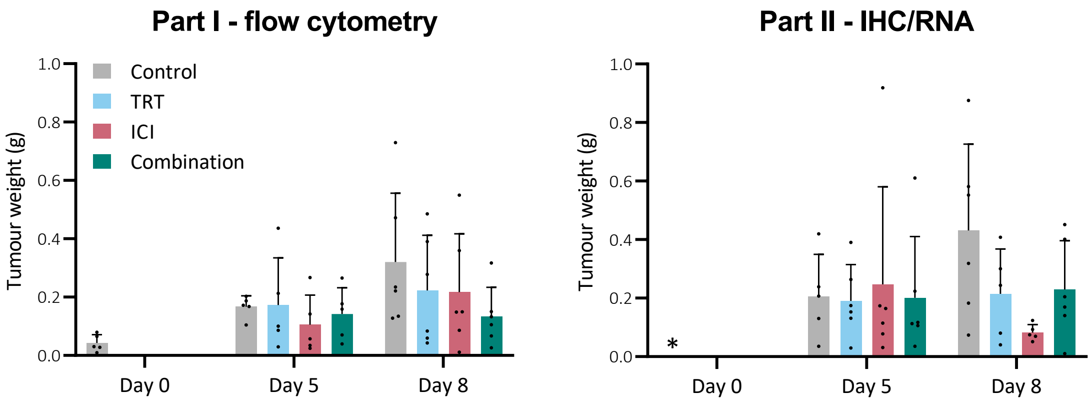


**Figure S6**. Renca-CAIX tumor weights after dissection, before processing for flow cytometry (left panel) or immunohistochemistry and RNA expression profiling (right panel). Data represents mean + SD of all mice (dots) per group. *Missing value.

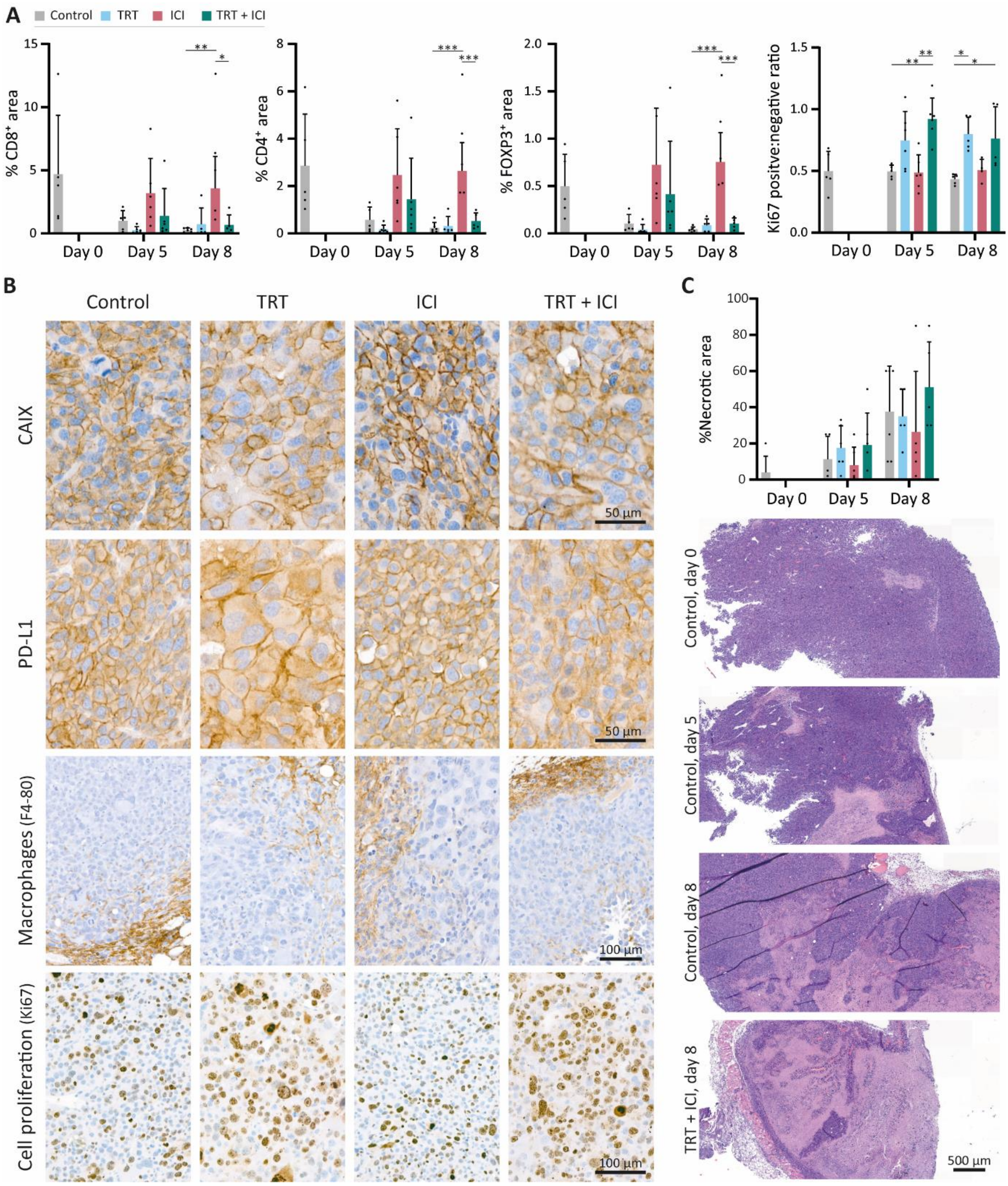


**Figure S7.** Renca-CAIX tumor-bearing balb/c mice (*n*=5-6/group) were treated with vehicle (control), [$^{177}$Lu]Lu-DOTA-hG250 (TRT), aPD-1 + aCTLA-4 (ICI) or TRT+ICI combination therapy and sacrificed on day 0, 5, or 8 after therapy. (**A**) Quantification of FFPE tumor sections stained for CD8, CD4, FOXP3, and Ki67. (**B**) Representative images of FFPE tumor sections (day 8 after treatment) stained for CAIX, PD-L1, F4-80 (macrophages) and Ki67 (cell proliferation). (**C**) Manual estimation of percentage of necrotic area with representative images of H&E stained tumor tissues from indicated groups. Data represent mean + SD of individual mice (dots) per group. Statistical differences between treatment groups on day 5 and day 8 were separately determined by one-way ANOVA analyses with Šídák's multiple comparisons test. (*p<0.033, **p<0.01, ***p<0.001).

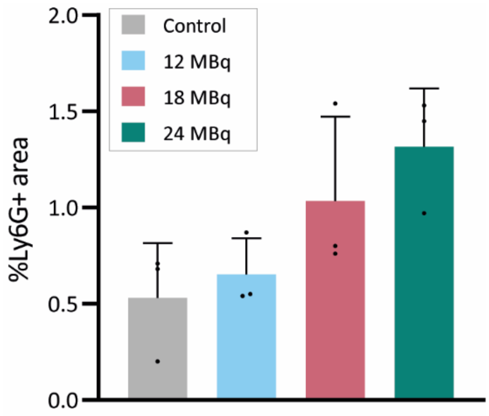


**Figure S8.** Neutrophil quantification of Ly6G-stained FFPE tumor sections, obtained from Renca-CAIX tumor-bearing mice one week after treatment with vehicle (control), 12 MBq, 18 MBq, or 24 MBq of [$^{177}$Lu]Lu-DOTA-hG250. Data represent mean percentage of Ly6G-positive area + SD of individual mice (dots, *n*=3/group).

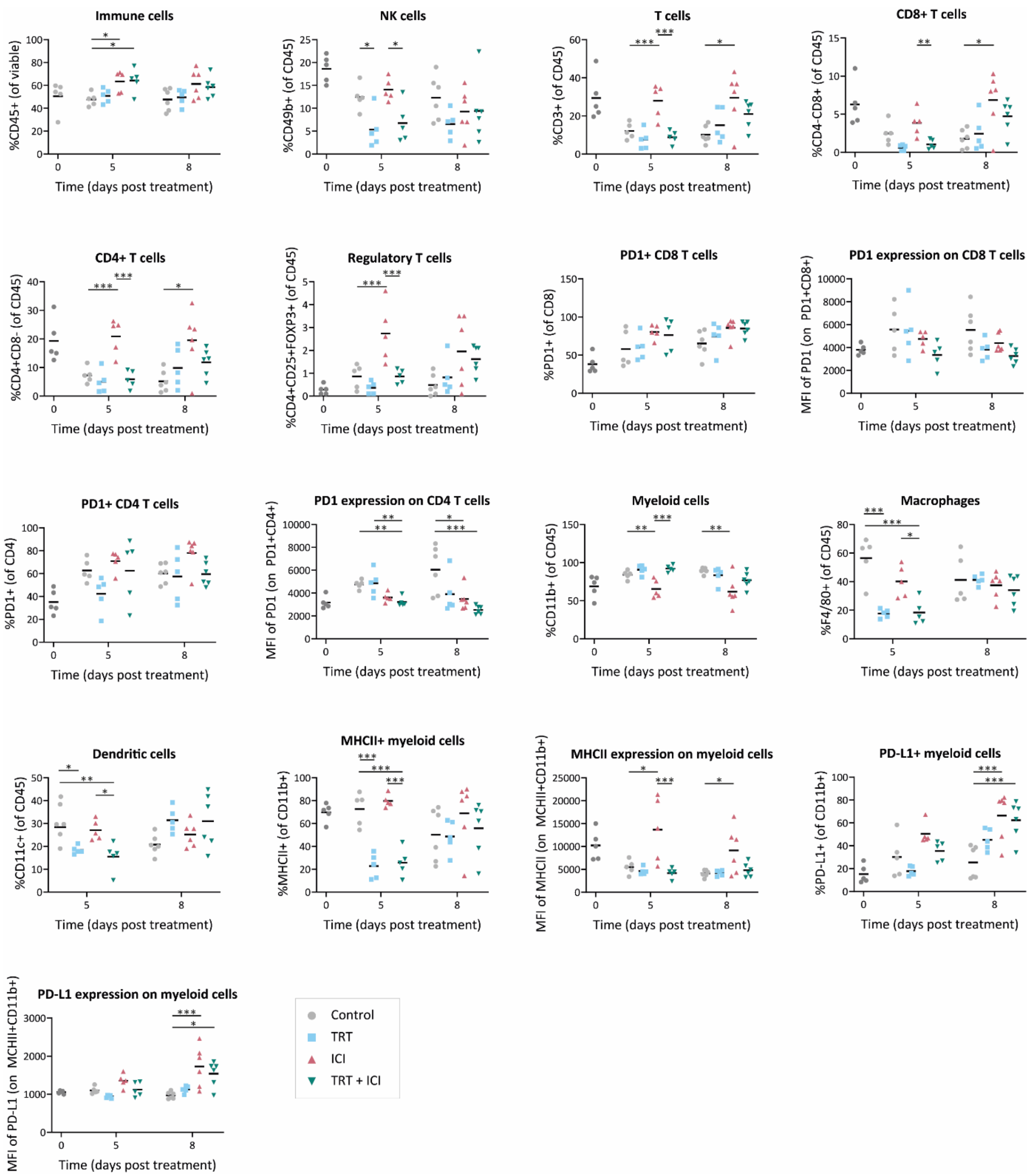


**Figure S9.** Flow cytometry analyses of tumor-infiltrating immune cells (CD45+), T cells (CD3+), cytotoxic T cells (CD3+CD4-CD8+), CD4 T cells (CD3+CD4+CD8-), Tregs (CD3+CD4+CD25+FOXP3+), myeloid cells (CD11b+), macrophages (CD11b+F4/80+), dendritic cells (CD11b+CD11c+), and NK cells (CD49b+) as percent of total immune cells and expression levels of PD1 (on T cells), MHCII and PD-L1 (on myeloid cells) as mean fluorescent intensity (MFI). Data represent mean (line) of individual mice (dots) per group. Statistical differences between treatment groups were determined by a mixed effects model (*p<0.05, **p<0.01, ***p<0.005).

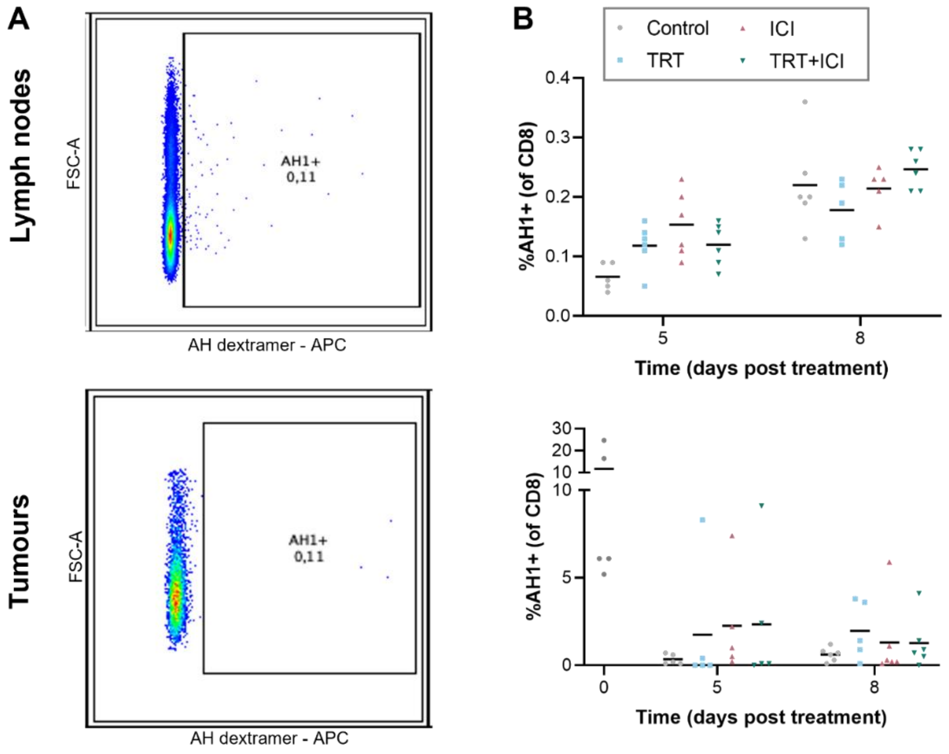


**Figure S10**. Flow cytometry analyses of AH antigen-specific CD8+ T cells in lymph nodes (upper panels) and tumors (lower panels). (**A**) Dot plots for AH dextramer-positive CD8+ T cells. (**B**) Graphs show mean (lines) percentage of AH1 dextramer+ CD8+ T cells of individual mice (dots) per group.

| Gene Set | TRT | ICI | Combi | Combi vs ICI |
|---|---|---|---|---|
| Angiogenesis | 1.48 | 2.1069 | 1.8109 | 2.0694 |
| Antigen Presentation | 1.8454 | 3.9659 | 3.2793 | 1.7747 |
| Apoptosis | 1.6529 | 2.7331 | 2.6219 | 1.9398 |
| Autophagy | 1.2285 | 1.8773 | 1.5517 | 2.3141 |
| Cell Proliferation | 1.6823 | 2.3143 | 1.9677 | 2.4061 |
| Costimulatory Signaling | 1.397 | 3.7231 | 2.7289 | 2.0797 |
| Cytokine and Chemokine Signaling | 1.6436 | 3.1789 | 2.7096 | 2.0705 |
| **Cytotoxicity** | 2.8059 | 3.9519 | 4.2399 | 1.4619 |
| DNA Damage Repair | 1.8271 | 2.1449 | 2.4099 | 2.3532 |
| Epigenetic Regulation | 1.4421 | 2.0479 | 1.6034 | 2.0132 |
| Hedgehog Signaling | 1.491 | 3.0089 | 2.8836 | 1.6479 |
| Hypoxia | 1.0134 | 2.2083 | 1.6494 | 2.2577 |
| Immune Cell Adhesion and Migration | 1.3906 | 3.3308 | 2.5899 | 2.0426 |
| **Interferon Signaling** | 2.8088 | 4.2379 | 4.2613 | 1.4533 |
| JAK-STAT Signaling | 1.7474 | 3.3626 | 2.7937 | 2.1537 |
| Lymphoid Compartment | 2.1289 | 4.0553 | 3.5324 | 1.8662 |
| **MAPK** | 1.5106 | 2.0861 | 2.0028 | 2.288 |
| Matrix Remodeling and Metastasis | 1.5608 | 2.379 | 2.0227 | 2.2003 |
| **Metabolic Stress** | 1.1921 | 2.1195 | 1.6895 | 2.2867 |
| Myeloid Compartment | 1.3788 | 2.4221 | 2.386 | 1.7699 |
| NF-kappaB Signaling | 1.6366 | 3.428 | 2.6181 | 1.462 |
| Notch Signaling | 1.6545 | 2.1706 | 2.3397 | 1.994 |
| PI3K-Akt | 1.5654 | 2.3366 | 2.0856 | 2.2845 |
| TGF-beta Signaling | 1.5515 | 2.2449 | 1.8506 | 1.7748 |
| Wnt Signaling | 1.6687 | 1.933 | 2.1293 | 1.9813 |

**Figure S11**. Undirected global significance scores for nanostring-annotated gene sets for pair-wise comparisons of treatment groups with control group and combination with ICI treatment groups, as determined by RNA expression nanostring analysis and obtained from the Rosalind Platform.

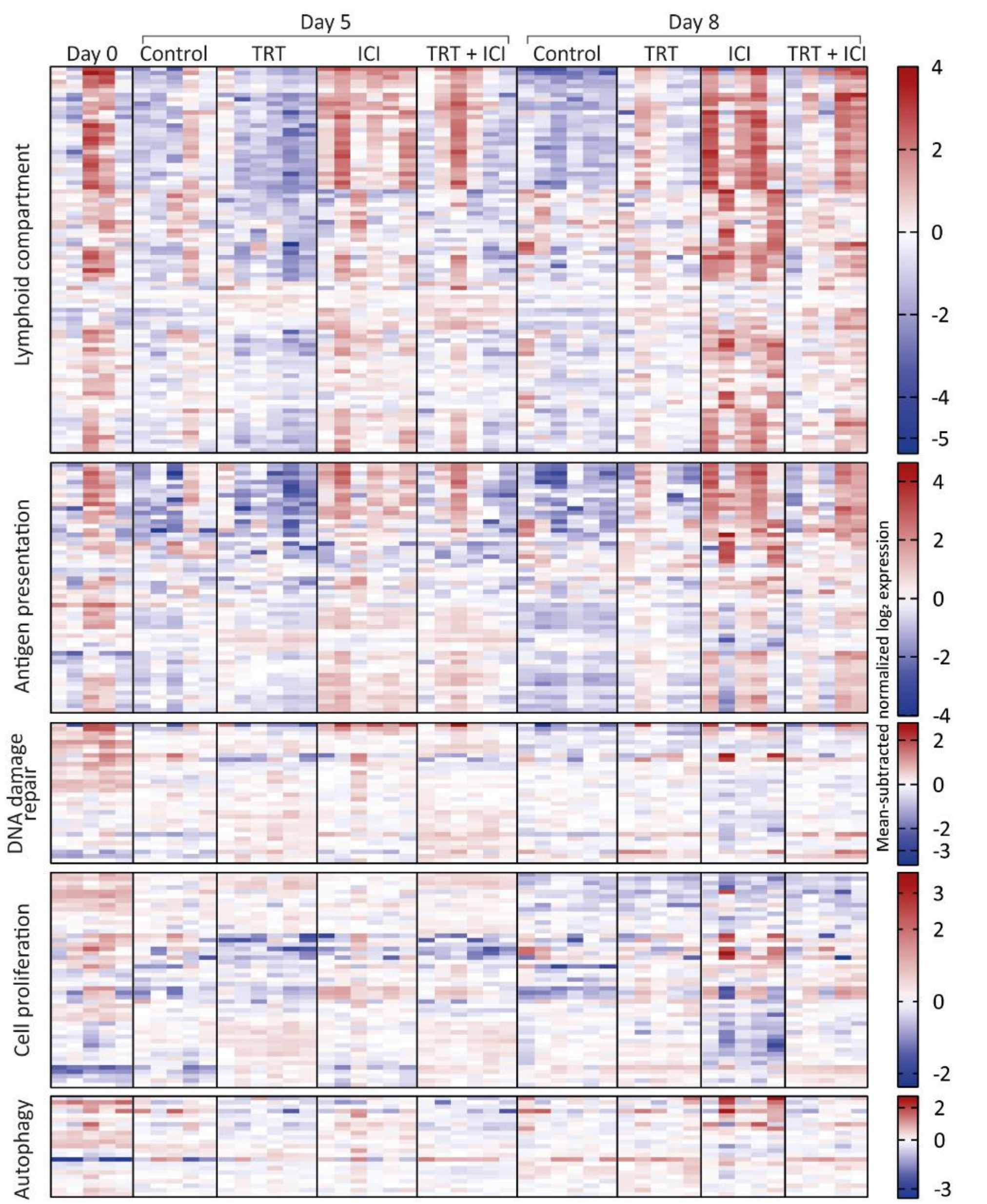


**Figure S12**. Nanostring gene expression analysis of tumors at 0, 5, or 8 days post treatment with vehicle (control), [$^{177}$Lu]Lu-DOTA-hG250 (TRT), aPD-1 + aCTLA-4 (ICI) or TRT + ICI combination therapy. Data represent heatmap of mean subtracted normalized log2 expression values for all samples of given gene sets.

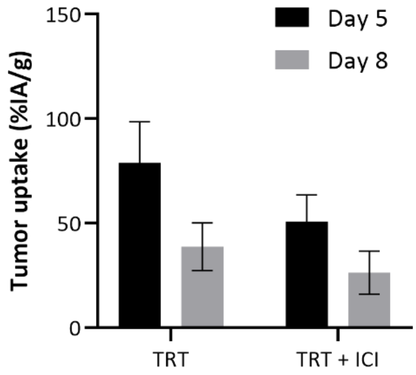


**Figure S13.** Renca-CAIX tumor uptake of [$^{177}$Lu]Lu-DOTA-hG250 (as %IA/g) in TME characterization experiment, 5 or 8 days after treatment with TRT only or TRT + ICI combination therapy.

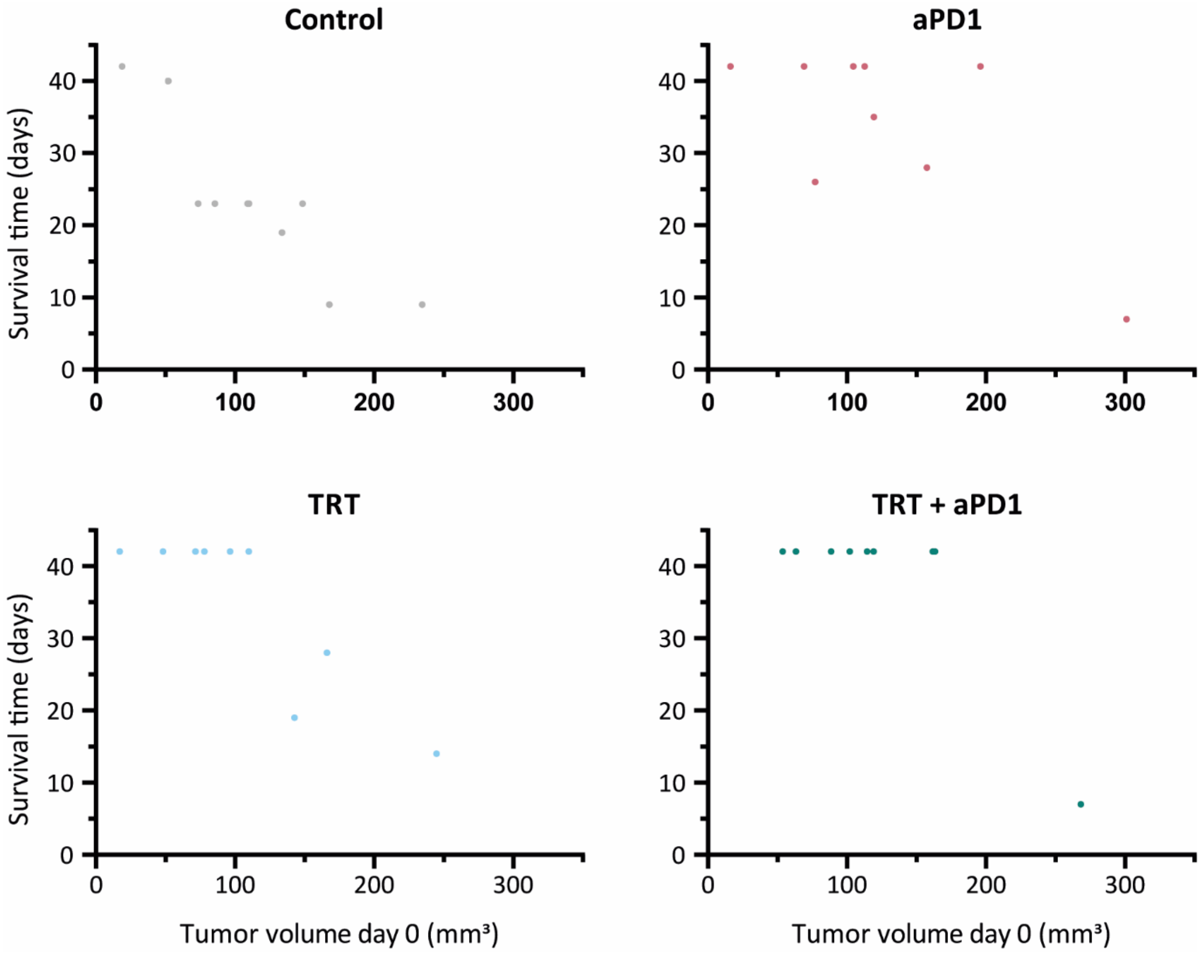


**Figure S14.** Treatment response, reported as survival time, of CT26-CAIX tumor-bearing mice after treatment with vehicle (control), aPD1, 4 MBq [$^{177}$Lu]Lu-DOTA-hG250 (TRT), or combined TRT+aPD1, plotted against tumor volume at start of treatment (day 0).

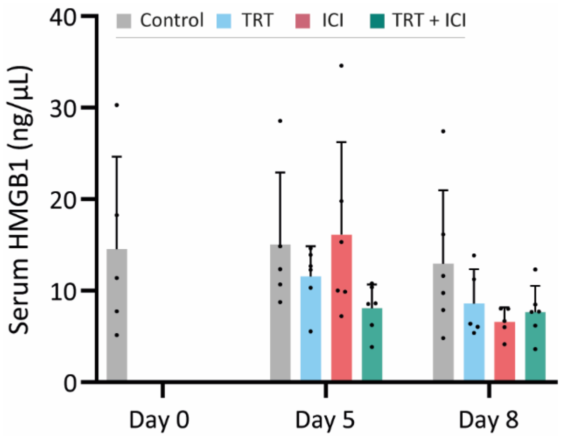


**Figure S15.** Serum concentration of HMGB1 of Renca-CAIX tumor-bearing mice after treatment with vehicle (control), aPD1+aCTLA4 (ICI), 4 MBq [$^{177}$Lu]Lu-DOTA-hG250 (TRT), or combined TRT+ICI. Data represent mean + SD. Statistical differences between treatment groups on day 5 and day 8 were separately determined by one-way ANOVA analyses with Šídák's multiple comparisons test and no statistical differences were found.
